# Claudin-low-like mouse mammary tumors show distinct transcriptomic patterns uncoupled from genomic drivers

**DOI:** 10.1101/557298

**Authors:** Christian Fougner, Helga Bergholtz, Raoul Kuiper, Jens Henrik Norum, Therese Sørlie

## Abstract

Claudin-low breast cancer is a molecular subtype associated with poor prognosis and without targeted treatment options. The claudin-low subtype is defined by certain biological characteristics, some of which may be clinically actionable, such as high immunogenicity. In mice, the medroxyprogesterone acetate (MPA) and 7,12-dimethylbenzanthracene (DMBA) induced mammary tumor model yields a heterogeneous set of tumors, a subset of which display claudin-low features. Neither the genomic characteristics of MPA/DMBA-induced claudin-low tumors, nor those of human claudin-low breast tumors, have been thoroughly explored.

The transcriptomic characteristics and subtypes of MPA/DMBA-induced mouse mammary tumors were determined using gene expression microarrays. Somatic mutations and copy number aberrations in MPA/DMBA-induced tumors were identified from whole exome sequencing data. A publicly available dataset was queried to explore the genomic characteristics of human claudin-low breast cancer and to validate findings in the murine tumors.

Half of MPA/DMBA-induced tumors showed a claudin-low-like subtype. All tumors carried mutations in known driver genes. While the specific genes carrying mutations varied between tumors, there was a consistent mutational signature with an overweight of T>A transversions in TG dinucleotides. Most tumors carried copy number aberrations with a potential oncogenic driver effect. Overall, several genomic events were observed recurrently, however none accurately delineated claudin-low-like tumors. Human claudin-low breast cancers carried a distinct set of genomic characteristics, in particular a relatively low burden of mutations and copy number aberrations. The gene expression characteristics of claudin-low-like MPA/DMBA-induced tumors accurately reflected those of human claudin-low tumors, including epithelial-mesenchymal transition phenotype, high level of immune activation and low degree of differentiation. There was an elevated expression of the immunosuppressive genes *PTGS2* (encoding COX-2) and *CD274* (encoding PD-L1) in human and murine claudin-low tumors. Our findings show that the claudin-low breast cancer subtype is not demarcated by specific genomic aberrations, but carries potentially targetable characteristics warranting further research.

**Author Summary:** Breast cancer is comprised of several distinct disease subtypes with different etiologies, prognoses and therapeutic targets. The claudin-low breast cancer subtype is relatively poorly understood, and no specific treatment exists targeting its unique characteristics. Animal models accurately representing human disease counterparts are vital for developing novel therapeutics, but for the claudin-low breast cancer subtype, no such uniform model exists. Here, we show that exposing mice to the carcinogen DMBA and the hormone MPA causes a diverse range of mammary tumors to grow, and half of these have a gene expression pattern similar to that seen in human claudin-low breast cancer. These tumors have numerous changes in their DNA, with clear differences between each tumor, however no specific DNA aberrations clearly demarcate the claudin-low subtype. We also analyzed human breast cancers and show that human claudin-low tumors have several clear patterns in their DNA aberrations, but no specific features accurately distinguish claudin-low from non-claudin-low breast cancer. Finally, we show that both human and murine claudin-low tumors express high levels of genes associated with suppression of immune response. In sum, we highlight claudin-low breast cancer as a clinically relevant subtype with a complex etiology, and with potential unexploited therapeutic targets.

## Introduction

The claudin-low subtype of breast cancer (BC) is a distinct disease entity associated with a relatively poor prognosis, and with an inadequately understood clinical significance [1–3]. It is characterized by low expression of tight junction and cell-cell adhesion genes, low degree of differentiation, epithelial-mesenchymal transition (EMT) phenotype and high level of immune cell infiltration [2]. The claudin-low subtype represents 7-14% of all breast cancers, and despite its unique biological features, there are no therapies specifically targeting the subtype [2–5]. While claudin-low tumors are found in several large scale studies, there is a paucity of information regarding their specific genomic characteristics [6–9]. Thus, significant gaps remain in the understanding of the biology of claudin-low tumors, and there is a need for further research to explore how their unique features may be therapeutically targeted.

Accurate preclinical models are vital for research into novel treatment options. Mouse mammary tumors may be induced through exposure to medroxyprogesterone acetate (MPA) and 7,12-dimethylbenzanthracene (DMBA) [10]. The tumors generated by this protocol are diverse, and a subset of these show similarities to the human claudin-low subtype [11,12]. A homogeneous primary *in vivo* model of claudin-low breast cancer does not currently exist [11]. While the mechanisms of MPA [10,13] and DMBA [14–17] have been described, there is still contention regarding the suitability of a chemically induced model of cancer for a disease that is not primarily caused by carcinogens in humans [18]. Evaluating the claudin-low subset of MPA/DMBA-induced tumors as a model for human disease is therefore an important step toward advancing preclinical research of claudin-low breast cancer.

In this study, we identified and comprehensively characterized claudin-low-like mouse mammary tumors generated by MPA/DMBA-induced carcinogenesis. Through genomic and transcriptomic analyses, we evaluated these tumors as a model for human claudin-low breast cancer and showed these tumors to be phenotypically accurate representations of their human counterparts. In parallel, we analyzed the previously unexplored genomic features of human claudin-low breast cancer. Our findings highlighted several features of claudin-low breast cancer with potential therapeutic implications, including a low tumor mutational burden, high expression of the immune checkpoint gene *CD274* (encoding PD-L1) and high expression of *PTGS2* (encoding cyclooxygenase-2).

## Results

### Gene expression subtyping reveals two distinct tumor clusters

We determined the murine transcriptomic subtypes of 17 MPA/DMBA-induced mammary tumors from 13 mice (S1 File) by performing a hierarchical clustering of gene expression data using the mouse intrinsic gene list [11]. This revealed nine murine subtypes in the cohort (Fig 1, Table 1, S2 File). The tumors separated into two distinct clusters. One cluster consisted of claudin-low^Ex^ and squamous-like^Ex^ tumors, both of which have been shown to resemble the human claudin-low subtype [11]; this is therefore referred to as the claudin-low-like cluster. The other cluster contained tumors from seven different subtypes and is referred to as the mixed cluster. In four instances, two tumors from different mammary glands were harvested from the same mouse. These were classified as different subtypes in all cases and are presumed to be distinct primary tumors. All normal mammary gland samples were classified as normal-like^Ex^, and clustered separately from the tumors.

**Fig 1.**
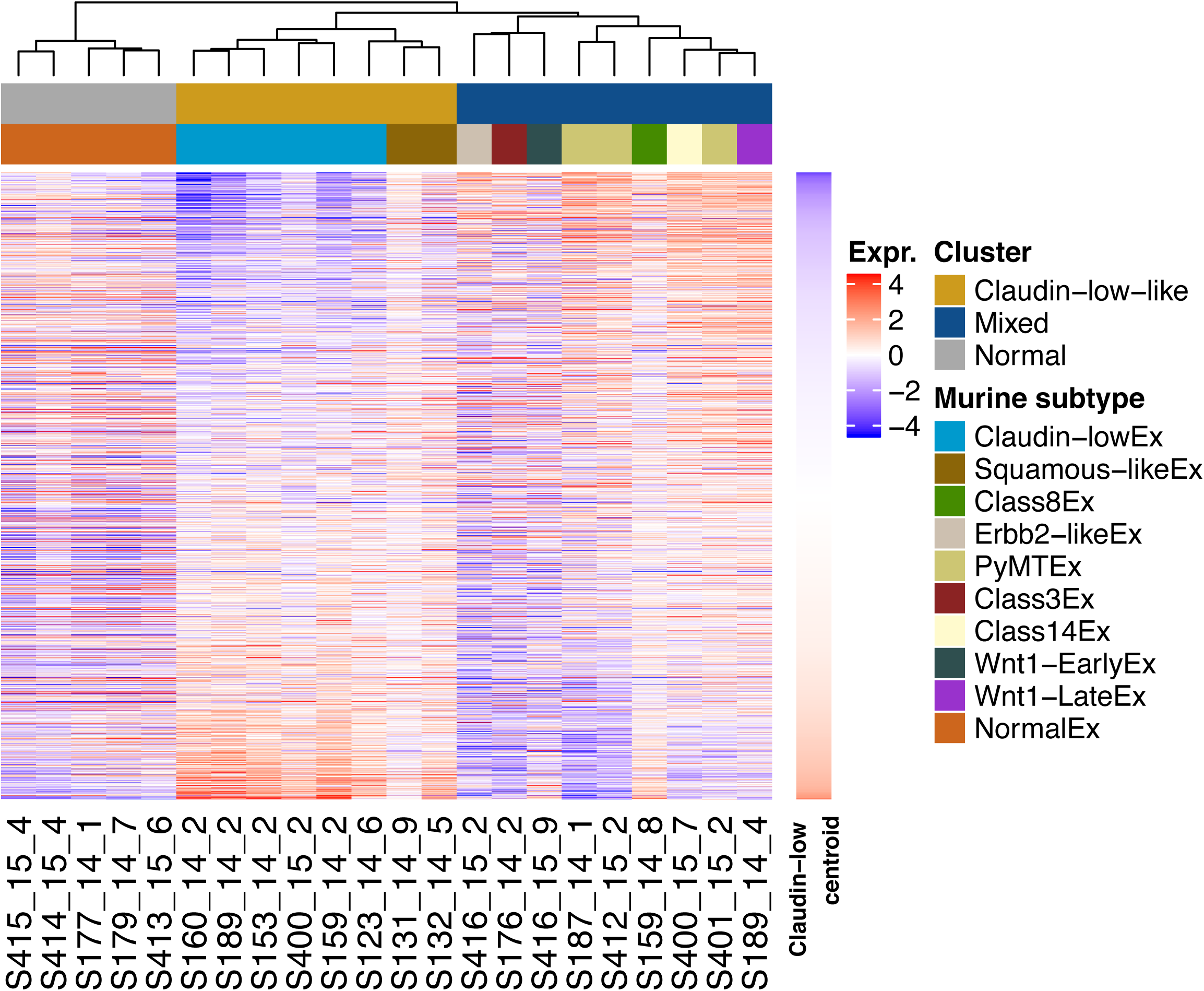
Gene expression-based subtypes in the MPA/DMBA-induced tumor cohort. Hierarchical clustering of MPA/DMBA-induced mouse mammary tumor gene expression levels revealed two distinct clusters, one containing claudin-low-like tumors and the other containing a transcriptomically heterogeneous set of tumors. Normal mouse mammary gland samples formed a separate cluster.

**Table 1.**
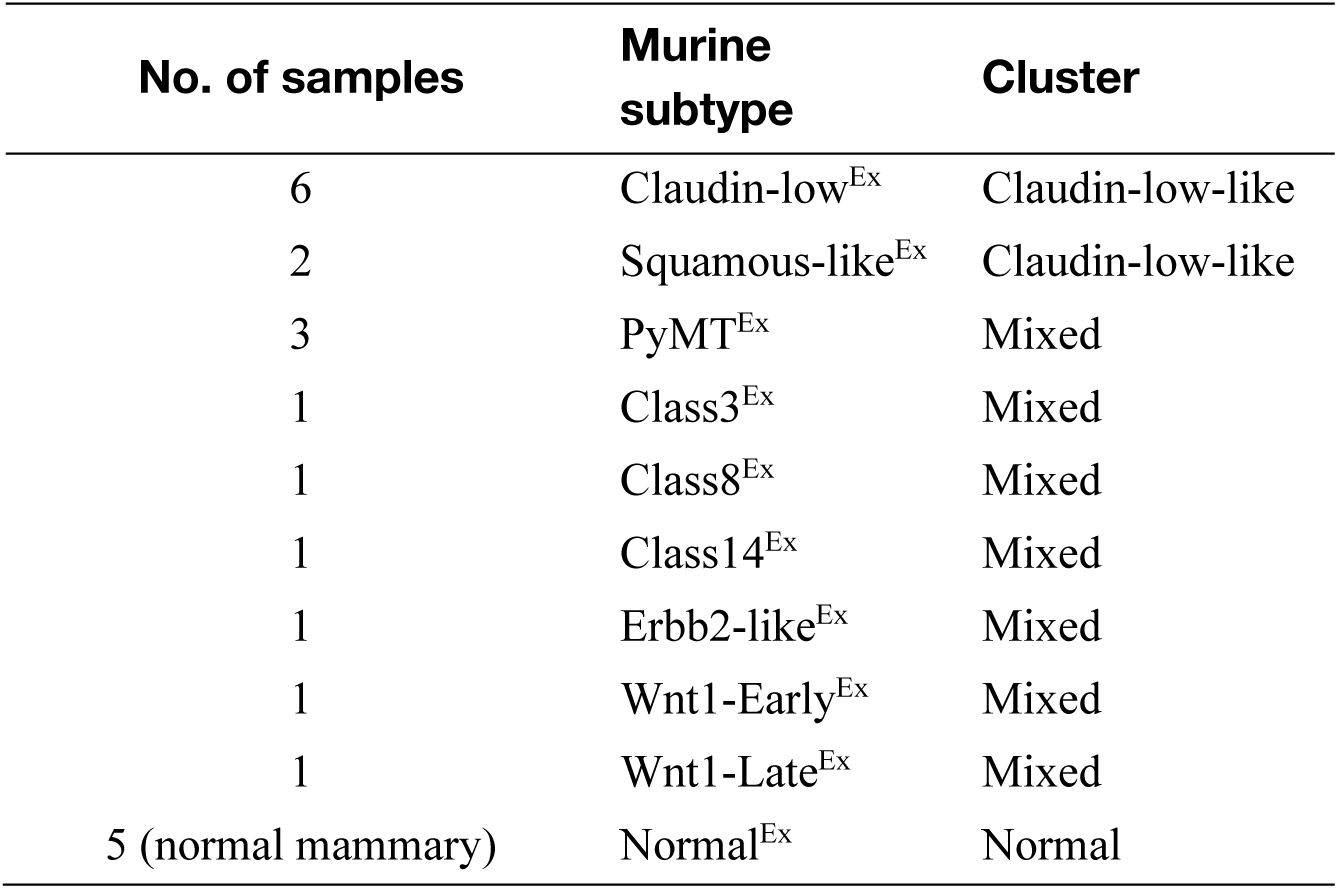
Subtype distribution of MPA/DMBA-induced tumors and normal mouse mammary gland tissue

| No. of samples | Murine subtype | Cluster |
| --- | --- | --- |
| 6 | Claudin-low<sup>Ex</sup> | Claudin-low-like |
| 2 | Squamous-like<sup>Ex</sup> | Claudin-low-like |
| 3 | PyMT<sup>Ex</sup> | Mixed |
| 1 | Class3<sup>Ex</sup> | Mixed |
| 1 | Class8<sup>Ex</sup> | Mixed |
| 1 | Class14<sup>Ex</sup> | Mixed |
| 1 | ErbB2-like<sup>Ex</sup> | Mixed |
| 1 | Wnt1-Early<sup>Ex</sup> | Mixed |
| 1 | Wnt1-Late<sup>Ex</sup> | Mixed |
| 5 (normal mammary) | Normal<sup>Ex</sup> | Normal |

Histopathological analysis corroborated the intertumor heterogeneity that was demonstrated by subtyping (S1 File). Five of the eight claudin-low-like tumors, including both squamous-like^Ex^ tumors, showed a squamous appearance, while no tumors in the mixed cluster displayed this histological phenotype (p = 0.009, Fisher’s exact test). There was also a higher frequency of claudin-low-like tumors showing marked neutrophil infiltration (p = 0.002, Fisher’s exact test) and displaying a marked or partial spindloid appearance (p = 0.050, Fisher’s exact test) compared to tumors in the mixed cluster.

### Mutations in MPA/DMBA-induced mammary tumors are independent of gene expression subtype

To determine the genetic characteristics of the tumors, we performed exome sequencing to a mean depth of 58, with 84% of bases being sequenced to a coverage of 20x or higher. We identified a mean of 589 mutations per tumor (range: 288 to 1795), corresponding to a mean mutation rate of 11.9 mutations per megabase (range: 5.8 to 36.2) (Fig 2B). This is substantially higher than the average 1.3 mutations per megabase found in human breast cancer [19]. There was no significant difference in mutational burden between the tumors in the claudin-low-like and the mixed cluster, and the only subtype specific trend was a particularly high mutational burden in the two squamous-like^Ex^ tumors (Fig 2B).

**Fig 2.**
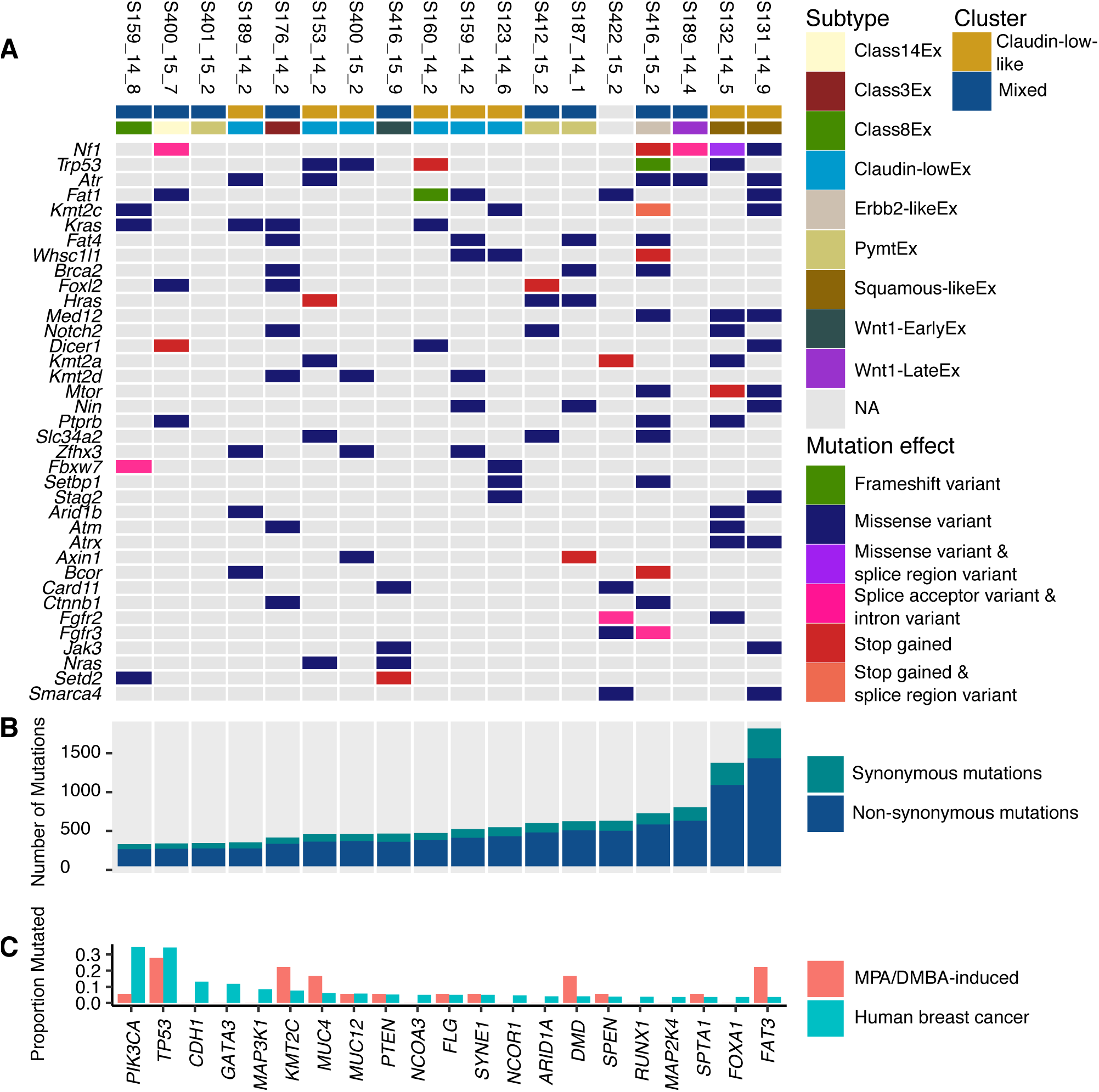
Somatic mutations in MPA/DMBA-induced mouse mammary tumors. **A** *Nf1, Trp53, Atr* and *Fat1* were the most frequently mutated driver genes in the MPA/DMBA-induced tumor cohort. No specific mutations accurately delineated the tumor clusters. **B** The MPA/DMBA-induced tumors carried between 288 and 1795 exonic mutations. No significant differences in mutational burden were found between the clusters, however a high mutational rate was observed in the two squamous-like^Ex^ tumors. **C** MPA/DMBA-induced tumors generally showed divergent mutational rates compared to human breast cancer in the genes most frequently mutated in human breast cancer. *TP53* mutations occurred at a similar rate in MPA/DMBA-induced tumors and human breast cancer.

All tumors carried mutations in driver genes defined by the COSMIC cancer gene census list [20], with a mean of 13.8 driver genes carrying mutations per tumor (range: 4 to 29) (Fig 2A). Several driver genes were recurrently mutated, including *Trp53, Kras*, and *Kmt2c* (S3 File), but no driver genes carried mutations at a significantly different rate between the two clusters. We did, however, identify two notable trends which did not reach statistical significance: an elevated rate of *Trp53* mutations in the claudin-low-like cluster (50% vs. 11%, p = 0.13, two-tailed Fisher’s exact test) and an elevated rate of *Zfhx3* mutations also in the claudin-low-like cluster (37.5% vs. 0%, p = 0.08, two-tailed Fisher’s exact test). No mutations were significantly associated with histological features.

### MPA/DMBA-induced tumors and human breast cancers display disparate gene mutational profiles

To narrow down potential driver mutations in the MPA/DMBA-induced tumors, we compared amino acid changes caused by mutations in driver genes to known amino acid changes in human cancers [20] (Table 2, S4 File). There were hotspot amino acid changes in all *Ras* genes, including *Kras* G12C, G13R, Q61H, *Hras* Q61L and *Nras* Q61L. In total, 8 of 18 tumors carried hotspot amino acid changes in *Ras* genes. There was one *Pik3ca* mutation in the cohort causing an H1047R amino acid change. This mutation is frequently found in human breast cancer and has previously been reported in DMBA-induced mouse mammary tumors [21].

**Table 2.**
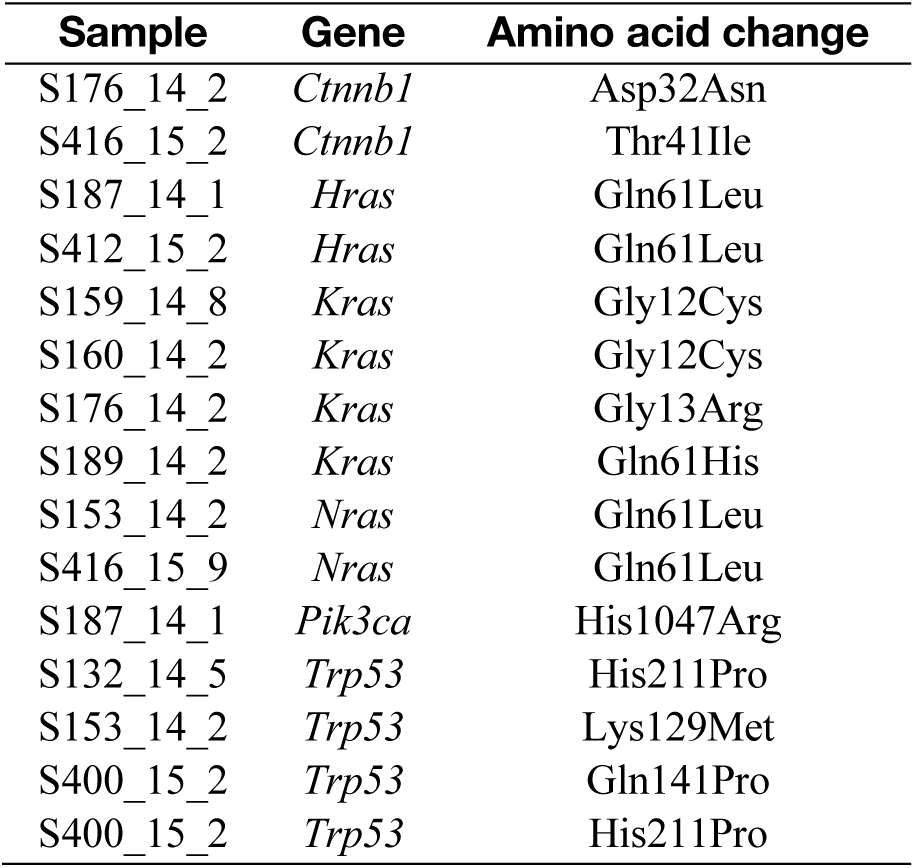
Selected hotspot mutations in MPA/DMBA-induced tumors

| Sample | Gene | Amino acid change |
| --- | --- | --- |
| S176_14_2 | *Cttnb1* | Asp32Asn |
| S416_15_2 | *Cttnb1* | Thr41Ile |
| S187_14_1 | *Hras* | Gln61Leu |
| S412_15_2 | *Hras* | Gln61Leu |
| S159_14_8 | *Kras* | Gly12Cys |
| S160_14_2 | *Kras* | Gly12Cys |
| S176_14_2 | *Kras* | Gly13Arg |
| S189_14_2 | *Kras* | Gln61His |
| S153_14_2 | *Nras* | Gln61Leu |
| S416_15_9 | *Nras* | Gln61Leu |
| S187_14_1 | *Pik3ca* | His1047Arg |
| S132_14_5 | *Trp53* | His211Pro |
| S153_14_2 | *Trp53* | Lys129Met |
| S400_15_2 | *Trp53* | Gln141Pro |
| S400_15_2 | *Trp53* | His211Pro |

There were marked disparities between the gene mutational profiles of human breast cancer [22] and MPA/DMBA-induced tumors (Fig 2C, S5 File). The two most frequently mutated genes in breast cancer are *PIK3CA* and *TP53*. While *TP53* showed comparable mutation rates between human breast cancer and MPA/DMBA-induced tumors (34% and 28%, respectively), *PIK3CA* mutation does not appear to be a common event in MPA/DMBA-induced tumors (35% in BC, 6% in MPA/DMBA). Several frequently mutated genes in breast cancer, such as *CDH1, GATA3* and *MAP3K1*, were not mutated in any MPA/DMBA-induced tumors. Conversely, many genes frequently mutated in MPA/DMBA-induced tumors, such as *ATR, FAT1* and *KRAS*, are rarely mutated in breast cancer.

### DMBA induces a characteristic mutational spectrum with a high frequency of T>A transversions in TG dinucleotides

To characterize the mutagenic profile of DMBA, we analyzed the mutational spectra of the MPA/DMBA-induced tumors. Mutations showed a majority of T>A transversions, which accounted for 63% of all mutations (S1A Fig). In their trinucleotide context, thymine mutations (T>N) were overrepresented in positions with a 3’ guanine nucleotide (S1B & S1C Fig, S6 File). This was statistically significant when compared to the proportion of thymine nucleotides in an NTG context in the mouse reference genome (p < 0.001 in all cases, two-tailed Wilcoxon rank-sum test). There was a similar overrepresentation of cytosine mutations in positions with a 3’ adenine. This was statistically significant for C>A and C>G mutations (p < 0.001), but not for C>T mutations (p = 0.089), when compared to the proportion of cytosine nucleotides in an NCA context in the mouse reference genome.

Mutation signature analysis revealed evidence of signatures 4, 6, 22, 24 and 25 [23] in the MPA/DMBA-induced tumors (S1D Fig). All tumors were associated with signature 22, while signatures 4 and 25 were found in 17 and 11 of the 18 tumors, respectively. Signatures 24 and 6 were only found in four and one tumor(s), respectively. Notably, none of the signatures found in MPA/DMBA-induced tumors have been associated with human breast cancer [23].

### MPA/DMBA-induced tumors have diverse copy number profiles

Breast cancer is largely driven by copy number aberrations (CNAs) [24], yet the copy number profiles of MPA/DMBA-induced mammary tumors have not previously been described. We found a mean of 1299 genes with CNA per tumor (range: 90 – 3057), of which a mean of 65% were amplifications. There was a tendency for claudin-low-like tumors to have a lower burden of CNAs, with a mean of 919 genes carrying CNA, compared to the mixed group of tumors, with a mean of 1637 genes carrying CNA (Fig 3A). This trend did however not reach statistical significance (p = 0.139, two-tailed Wilcoxon rank-sum test).

**Fig 3.**
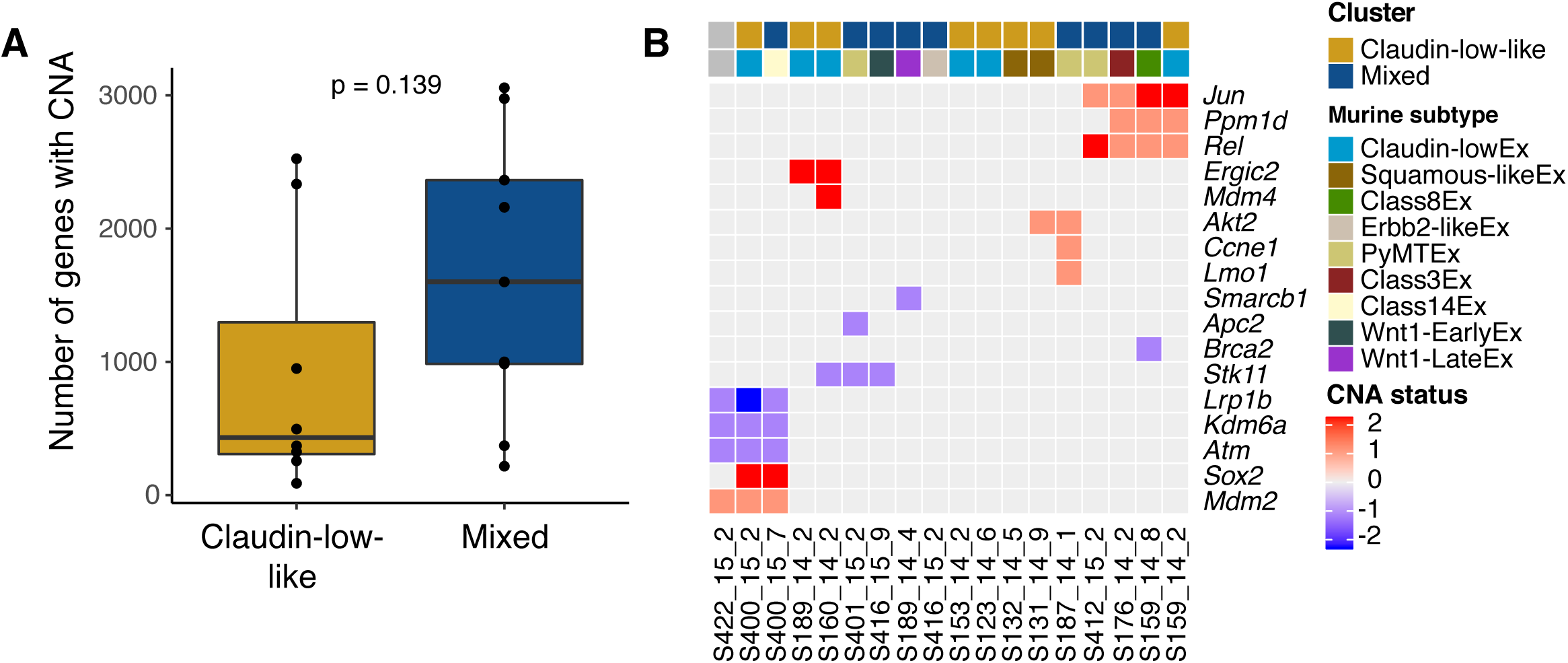
Copy number aberrations in MPA/DMBA-induced mouse mammary tumors. **A** There was a trend toward a lower number of genes with copy number aberrations in the claudin-low-like cluster. **B** Copy number aberrations implicated in cancer were found in 14 of 18 MPA/DMBA-induced tumors. Two tumor sets (*S422_15_2, S400_15_2* and *S400_15_7*, and *S412_15_2, S176_14_2, S159_14_8* and *S159_14_2*) showed remarkably similar CNA profiles, but displayed different gene expression subtypes. CNA status of −2 is a homozygous deletion, CNA status of −1 is a heterozygous deletion, CNA status of 0 is copy number neutral, CNA status of 1 is a single copy amplification, and CNA status of 2 is a multi-copy amplification.

To determine CNAs in the MPA/DMBA-induced tumors with a potential oncogenic driver effect, we identified amplifications and deletions known to be associated with cancer [20] (Fig 3B). We found that 14 of the 18 tumors carried potential driver CNAs (range: 0 to 4, mean: 2.6). Three of the four tumors not carrying potential driver CNAs were claudin-low-like. There was however no statistically significant difference in the number of potential driver CNAs between the clusters. Several genes had recurrent CNAs, but none occurred at a statistically significant different rate in one cluster versus the other.

Only two of the CNA events identified in MPA/DMBA-induced tumors occur at a notable rate in human breast cancer; *Mdm4* is amplified in 25% and *Ppm1d* is amplified in 10% of human BC [6,7].

We observed two sets of tumors carrying remarkably similar CNA profiles (Fig 3B). None of the tumors in these two sets displayed the same murine subtype as any other tumor within the same set.

### The human claudin-low breast cancer genome is characterized by a low mutational burden, frequent *TP53* mutations and a low rate of CNA

Little has been published specifically describing the genomic characteristics of human claudin-low breast cancer. We therefore analyzed the 218 claudin-low tumors found in the METABRIC dataset, for which DNA sequence data from 173 genes and whole genome copy number data is available [6,7].

Claudin-low tumors, with a mean of 4.7 mutations per tumor, carried relatively few mutations compared to all other tumors, with a mean of 7.3 mutations per tumor (p < 0.001, two-tailed Wilcoxon rank-sum test) (Fig 4A). Claudin-low tumors share several characteristics with basal-like tumors and are often classified as such by the PAM50 assay [6,7,25]; however, basal-like tumors showed a significantly higher mutational burden than claudin-low tumors (mean: 8.08 mutations per tumor, p < 0.001, two-tailed Wilcoxon rank-sum test).

**Fig 4.**
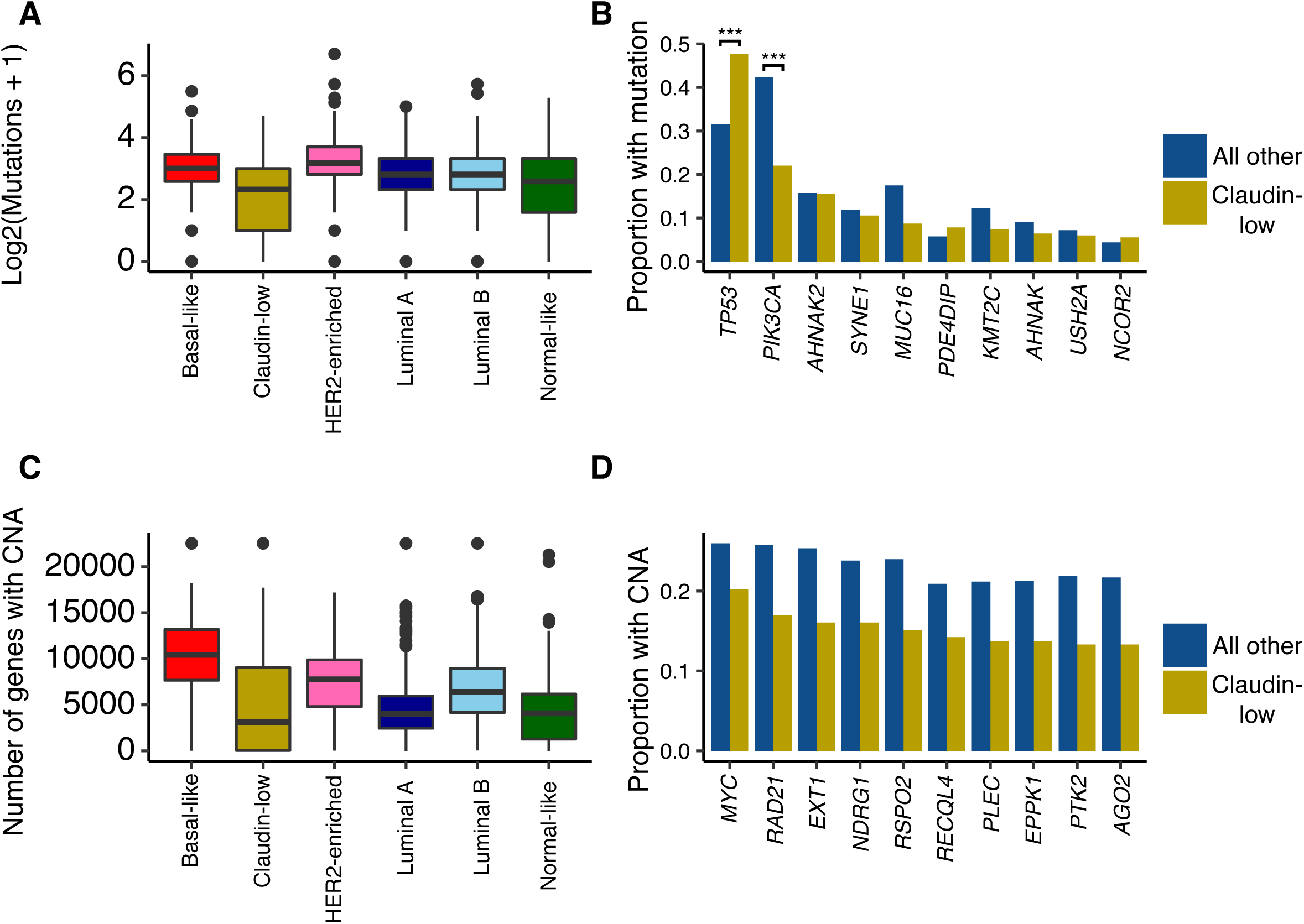
Somatic mutations and copy number aberrations in human claudin-low breast cancer. **A** Claudin-low breast cancer was the subtype with the lowest mutational burden. Number of mutations displayed as log_2_(mutations + 1). **B** *TP53* and *PIK3CA* were the most frequently mutated genes in human breast cancer. Claudin-low tumors carried *TP53* and *PIK3CA* mutations at significantly higher and lower rates, respectively, compared to non-claudin-low breast tumors. *** = p < 0.001. **C** Claudin-low tumors carried relatively few CNAs compared to non-claudin-low tumors. **D** The ten genes which were most frequently affected by CNA in claudin-low tumors were all found to be copy number aberrant at a higher frequency in non-claudin-low tumors. *MYC* amplification is the most common CNA event in claudin-low breast cancer.

There was a high degree of overlap between the genes most frequently mutated in claudin-low breast cancers and the genes most frequently mutated in all other breast cancers (Fig 4B). Most of these genes carried mutations at similar rates between claudin-low and non-claudin-low tumors, albeit with a tendency towards a slightly lower rate in claudin-low tumors. There were however two notable differences in mutational frequency: a significantly higher rate of *TP53* mutations and a significantly lower rate of *PIK3CA* mutations in claudin-low tumors compared to other tumors. Similarly, basal-like tumors also carried a high frequency of *TP53* mutations and a low frequency of *PIK3CA* mutations [7,22].

Human claudin-low breast tumors carried significantly fewer genes with copy number aberration (mean: 4879) compared to all other tumors (mean: 6247; p < 0.001, two-tailed Wilcoxon rank-sum test) (Fig 4C). This difference was also marked when comparing claudin-low tumors with basal like tumors (mean: 10175 genes per tumor; p < 0.001, two-tailed Wilcoxon rank-sum test).

By gene, the most frequent copy number event in claudin-low breast cancer was *MYC* amplification, found in 20% of cases (Fig 4D). In comparison, this event was found in 26% of all other breast tumors. The ten most frequently amplified genes in claudin-low breast cancer were all located at chromosomal position 8q24, a region also frequently amplified in basal-like breast cancers [6,7].

### Claudin-low-like MPA/DMBA-induced mammary tumors accurately reflect the gene expression characteristics of their human counterpart

We explored several established gene expression features of the claudin-low subtype and found that MPA/DMBA-induced claudin-low-like tumors accurately mirrored their human counterpart. Specifically, claudin-low-like tumors had low expression of genes involved in cell-cell adhesion, low expression of luminal genes, and high expression of genes related to EMT (Fig 5A, S2 Fig). Claudin-low-like tumors also showed a markedly lower degree of differentiation compared to tumors in the mixed cluster. In particular, the claudin-low-like cluster expressed significantly higher and lower levels of *Cd44* and *Cd24a*, respectively, indicating a stem cell-like phenotype in these tumors [25,26] (S3 Fig). There was no significant difference in the expression of proliferation-related genes between the two clusters. Vascular content-related genes were expressed at a significantly higher level in claudin-low-like tumors compared to the tumors in the mixed cluster (S2 Fig), indicating a higher degree of neoangiogenesis in these tumors.

**Fig 5.**
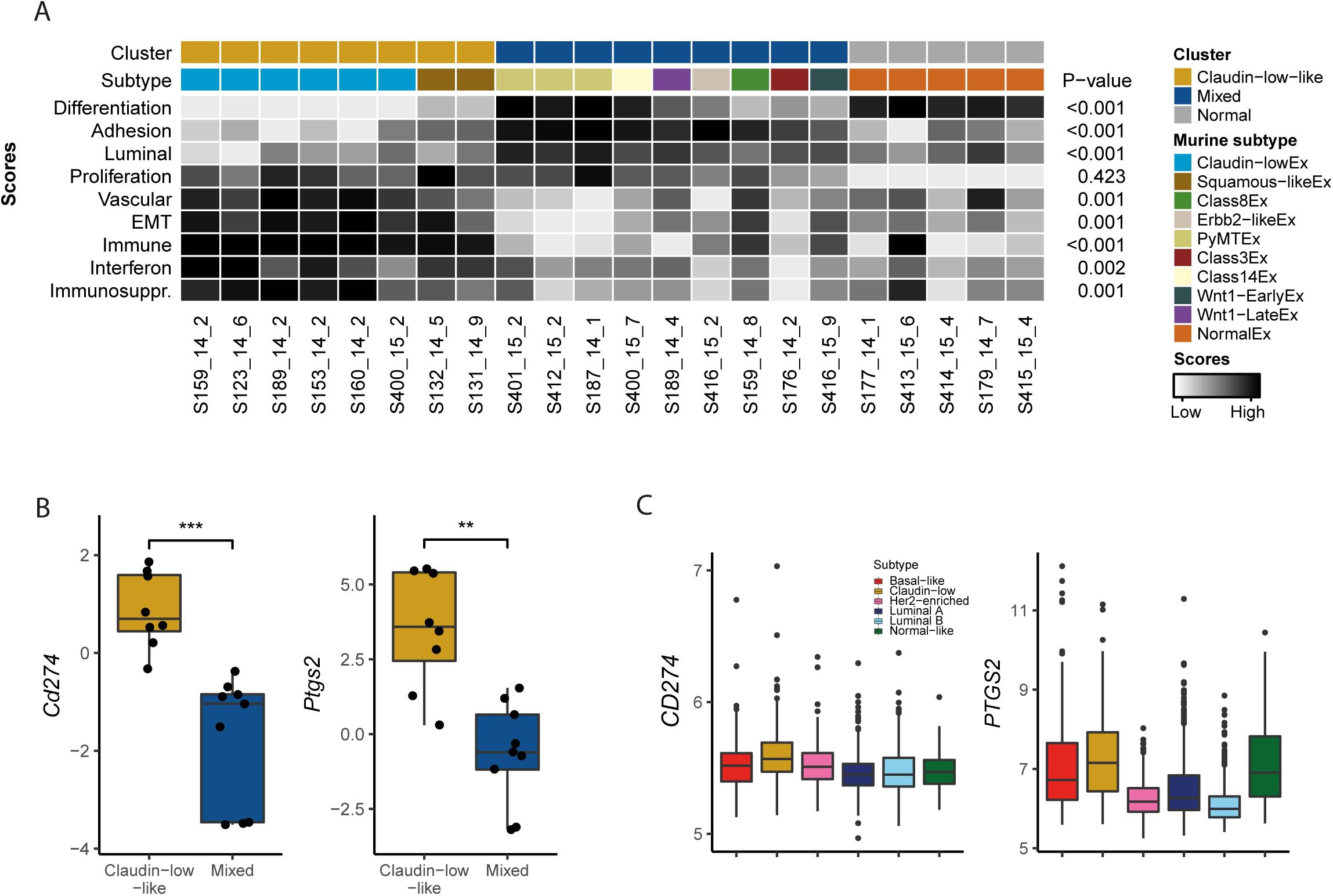
Gene expression characteristics of claudin-low-like MPA/DMBA-induced tumors and human claudin-low breast cancers. **A** MPA/DMBA-induced claudin-low-like tumors recapitulated the gene expression characteristics of the claudin-low subtype as evidenced by the expression levels of relevant gene signatures. P-values are calculated for the claudin-low-like tumors versus tumors in the mixed cluster. **B** *Cd274* and *Ptgs2* are expressed at significantly higher levels in the claudin-low-like tumors than in the mixed cluster tumors. **C** Claudin-low is the breast cancer subtype with the highest expression of *CD274* and *PTGS2*.

Immune cell admixture was significantly higher in the claudin-low-like tumors compared to tumors in the mixed cluster (p < 0.001, two-tailed Wilcoxon rank-sum test) and compared to normal mammary gland samples (p = 0.006). We also found higher expression of genes related to immunosuppression and interferons in the claudin-low-like cluster compared to both the mixed cluster and normal mammary gland samples. High immune cell infiltration combined with high expression of type 1 interferon-related and immunosuppressive genes are characteristics of tumors that may respond to immunotherapeutics [27,28].

We identified a significantly elevated expression of two potentially actionable genes related to immunosuppression in the claudin-low-like tumors: the immune checkpoint encoding gene *Cd274* and the cyclooxygenase encoding gene *Ptgs2* (Fig 5B). These features were also characteristic of human claudin-low tumors in the METABRIC cohort [6,7], which showed significantly higher expression levels of both *PTGS2* and *CD274* compared to non-claudin-low breast tumors (p < 0.001 for both, two-tailed Wilcoxon rank-sum test), and compared specifically to basal-like tumors (p = 0.004 and p < 0.001, respectively) (Fig 5C). These characteristics may indicate a susceptibility to immune checkpoint inhibitors and cyclooxygenase inhibitors in human claudin-low breast cancer [29,30].

## Discussion

In this study, we have performed a comprehensive analysis of mutations, copy number aberrations and gene expression characteristics of MPA/DMBA-induced tumors. We found marked inter-tumor heterogeneity, and showed that half of the tumors displayed a claudin-low-like phenotype, in line with a previous report [11]. Our findings demonstrate that these tumors provide a transcriptomically accurate representation of human claudin-low breast tumors, reflecting key features such as an EMT phenotype, high level of immune infiltration and a low degree of differentiation.

MPA/DMBA-induced tumors carried a mutational burden multiple times that of human breast cancer, a high frequency of activating *Ras-*mutations and a characteristic mutational spectrum. The specific genes carrying mutations varied widely between tumors, however all tumors had a consistent mutational signature. This indicates that the dominant mutational process in these tumors is DMBA-induced mutagenesis, and not aberrations occurring after tumor initiation, as a result of e.g. disrupted DNA-repair. Copy number aberrations in MPA/DMBA-induced tumors have not previously been explored, and we show here that most tumors carry potential driver CNAs. However, while we noted several genomic trends, such as a higher rate of *Trp53* mutation and a lower burden of CNA in MPA/DMBA-induced claudin-low-like tumors, no individual genomic features accurately delineated the two gene expression-based tumor clusters. Further, several tumors carried similar sets of mutations and/or CNAs but displayed different subtypes. This suggests that no specific genomic event determines tumor subtype, and that other etiological models may be more appropriate, such as different cells-of-origin [31] or microenvironmental factors [32]. This finding concurs with recent reports showing that transgenic mouse mammary tumors display histological and transcriptomic phenotypes largely uncoupled from their underlying driver mutations [33,34]. One possible model for MPA/DMBA-induced tumorigenesis is therefore as follows: first, MPA induces a RANK-l mediated mammary gland proliferation [10,13]. DMBA then induces mutations in mammary cells in a pattern as elucidated by our mutation signature analysis, predominantly in TG and CA dinucleotides, stochastically distributed throughout the genome. The tumor is initiated when one or more driver mutations occur, for example *Trp53* or *Ras*-mutation, with the tumor phenotype, however, determined by non-genomic factors. The biochemical mechanism of DMBA-induced mutagenesis has been described [14,15], whereas no causal mechanism for DMBA-induced copy number aberration is known; it is therefore likely that CNAs arise after tumor initiation.

Previous genomic analyses which included human claudin-low breast tumors have either not included specific analyses of the subtype [6,7], included few samples [3], or have been restricted to the triple-negative [35,36] or metaplastic [37] subsets of claudin-low tumors. We show here that human claudin-low tumors are characterized by a low number of mutations and a low burden of CNAs. This finding is surprising, given the apparent inverse correlation between CNA and mutational burden in cancer [24], and indicates that the claudin-low subtype is relatively genomically stable compared to other breast cancers. We also find similarities in genomic characteristics between claudin-low tumors and basal-like tumors, in particular a high frequency of *TP53* mutations, a low frequency of *PIK3CA* mutations, and 8q24 amplifications as a common event. While the transcriptomic similarity between these two subtypes is established [25], these findings illustrate that there are also marked genomic similarities between claudin-low and basal breast cancer, albeit with a lower burden of genomic aberrations in claudin-low tumors.

Claudin-low tumors show high expression of immune-related genes and a high level of immune cell infiltration [3,25,38]. However, claudin-low tumors also express high levels of immunosuppressive genes. In MPA/DMBA-induced claudin-low-like tumors, we observed an elevated expression of two particularly notable genes involved in immunosuppression: *Ptgs2* (encoding COX-2) and *Cd274* (encoding PD-L1). This observation was consistent in human claudin-low breast cancer. COX-2 may be implicated in cancer development through several mechanisms: reducing apoptosis, increasing epithelial cell proliferation, promoting angiogenesis, increasing invasiveness of tumor cells and immunosuppression [39–41]. COX-2 may also be involved in vasculogenic mimicry, a process in which epithelial tumor cells form vascular channel-like structures without participation of endothelial cells, allowing nutrients to reach tumor cells without the need for neoangiogenesis [42]. Vasculogenic mimicry has previously been shown to occur in claudin-low tumors [43]. COX-2 and PD-L1 are clinically actionable through the use of COX-inhibitors [29] and checkpoint inhibitors [44], respectively. Further research into the potential use of checkpoint inhibitors and COX-inhibitors in claudin-low breast cancer is warranted, with promising future avenues including combinatorial Treg-depletion [38].

In summary, we have found that claudin-low-like MPA/DMBA-induced mouse mammary tumors are a transcriptomically accurate model for human claudin-low breast cancer. We did not find strong evidence that claudin-low-like MPA/DMBA-induced tumors are delineated by any specific genomic features, however the relatively small number of samples included in this study may have obscured possible associations. By analyzing publicly available data, we showed that human claudin-low breast cancer is a relatively genomically stable subtype. There is a high expression of genes related to immunosuppression in claudin-low breast cancers, a feature which is evident in claudin-low-like MPA/DMBA-induced tumors. Our observations suggest immunosuppression as a potential therapeutic target in claudin-low breast cancer and indicate MPA/DMBA-induced claudin-low-like tumors as an appropriate model for continued research.

## Material and Methods

### Ethics statement

The Norwegian Food Safety Authority approved all experiments in advance of their implementation (FOTS ID #4385). All mice were bred and maintained within a specific pathogen free (SPF) barrier facility according to local and national regulations, with food and water *ad libitum*. For invasive procedures, animals were anaesthetized with sevoflurane gas. Animals were euthanized by cervical dislocation under anesthesia with sevoflurane gas.

### Mouse strains and tumor induction

Animals from an Lgr5-creERT2-EGFP;Rb/C transgenic mouse strain on a pure FVB/N background were purchased from Jackson Laboratory (www.jax.org), kindly gifted from Rune Toftgård (*Karolinska Institutet*, Sweden) and locally bred. The transgenes are considered biologically inert, and offspring from all potential genotypes were used (wildtype, single transgene and double transgene). Female mice were treated with medroxyprogesterone acetate (MPA) and 7,12-dimethylbenzanthracene (DMBA) in accordance with established protocol [10]. In brief: 90-day release MPA pellets (50 mg/pellet, Innovative Research of America cat.# NP-161) were implanted subcutaneously at 6 and 19 weeks after birth. 1 mg DMBA (Sigma Aldrich cat.# D3254) dissolved in corn oil (Sigma Aldrich cat.# C8267) was administered by oral gavage at 9, 10, 12 and 13 weeks after birth. Tumor growth was regularly monitored by manual palpation and measured by caliper. Tumor volume was estimated using the following formula: volume = (width^2^* length)/2. When the tumors reached the maximum allowed size of 1000mm^3^, or at the age of 32 weeks, tissue was collected at necropsy and fixed in 4% paraformaldehyde (PFA) or snap frozen and stored at −80 °C. 18 tumors from 14 mice, of which four mice carried two mammary tumors, were subject to genomic and transcriptomic analyses. Six normal mammary glands collected from mice not undergoing MPA/DMBA-treatment were included as controls. Mouse features and histopathological tumor features, can be found in S1 File.

### Histopathology and immunohistochemistry

Mouse tissue was fixed overnight in 4% PFA, routinely processed and paraffin embedded. Formalin-fixed paraffin-embedded tissue was sectioned and stained with hematoxylin and eosin (HE). HE-stained tissue was classified by a certified veterinary pathologist.

Immunohistochemical staining was performed as previously described [45] with primary antibodies against K5 (Covance cat.# PRB-160P), K18 (Progen cat.# 61028), Ki67 (Novocastra cat.# NCL-Ki67p), ER*α* (Millipore cat.# 06-935), PR (Abcam cat.# ab131486) and Her2/Erbb2 (Millipore cat.# 06-562).

### DNA and RNA isolation

DNA isolation for exome sequencing was carried out at Theragen Etex Bio Institute (Seoul, South Korea). DNA was isolated using QIAamp DNA Mini Kit (Qiagen cat.# 51306) per manufacturer’s protocol. DNA from two samples (*S159_14_11* and *S176_14_11*) was isolated using CTAB Extraction Solution (Biosesang cat.# C2007) per manufacturer’s protocol. DNA integrity was assessed by electrophoresis and concentration was determined using the Nanodrop ND-1000 spectrophotometer (Thermo Scientific cat.# ND-1000) and Qubit fluorometer (Thermo Scientific cat.# Q33226). Total RNA and DNA isolation for gene expression microarrays was carried out using the QIAcube system (Qiagen cat.# 9001292) with the AllPrep DNA/RNA Universal Kit (Qiagen cat.# 80224) according to the protocol provided by the supplier, with 30mg tissue as input. The tissue was manually minced with a scalpel on ice followed by lysis and homogenization using TissueLyzer LT (Qiagen cat.# 85600) and Qiashredder (Qiagen cat.# 79654), respectively. Nucleic acid concentrations were measured by NanoDrop ND-1000 spectrophotometer and RNA integrity was analyzed using Agilent 2100 Bioanalyzer (Agilent Technologies cat.# G2939BA).

### Gene expression microarrays

Gene expression profiling was performed using RNA isolated from eighteen snap frozen MPA/DMBA-induced tumors and six normal/untreated mouse mammary gland samples. Whole genome expression data was obtained using Agilent Sureprint G3 Mouse Gene Expression 8×60K microarrays (Agilent Technologies cat.# G4852B) with Low Input Quick Amp Labeling protocol (Agilent Technologies cat.# 5190-2331) and the Cy3 fluorophore. 40ng RNA was used for input. Microarrays were scanned using an Agilent SureScan Microarray Scanner (Agilent Technologies cat.# G4900DA) and data was extracted using Agilent Feature Extraction software. One tumor sample (*S422_15_2*) failed quality control and was excluded from further gene expression analyses.

### Gene expression analyses

Gene expression data was analyzed using Qlucore Omics Explorer 3.2 (Qlucore AB) and R version 3.3.2 [46]. Gene expression values were quantile normalized and probes with a standard deviation of less than 2.8% of the largest observed standard deviation were filtered out. For genes represented by more than one probe, mean expression values were calculated to obtain one gene expression value per gene. Principal component analysis was performed to assess data quality and one normal mammary gland sample (*S178_14_2*) was identified as an outlier and removed from further analysis. Murine subtypes were determined by first calculating centroids for each subtype using the original data from the mouse intrinsic gene list [11], followed by calculating Spearman correlation for every sample to each of the subtype centroids. The subtype with the highest correlation coefficient was assigned as the sample’s subtype.

Unsupervised hierarchical clustering was performed using average linkage and Spearman correlation as the distance metric. Immune cell infiltration was inferred using ESTIMATE [47]. Scores for gene signatures relevant to the claudin-low subtype (adhesion, EMT, luminalness, proliferation, vascular content, immunosuppression and interferons [25,43,48–50]) were calculated using a standard (Z) score approach: for every gene in each signature, a standardized expression value was calculated by subtracting the mean across all samples, then dividing by the standard deviation. Calculation of the mean of the standardized expression values across all genes in the signature yielded the score. Gene lists included in the different signatures are found in S7 File. The degree of differentiation was calculated using a differentiation predictor [25]. Two-tailed Wilcoxon rank-sum tests were used for statistical testing of differences in scores between two groups.

### Whole exome sequencing

Whole exome sequencing was carried out at Theragen Etex Bio Institute. Library preparation and target enrichment was carried out using the SureSelect XT Mouse All Exon Kit (Agilent cat.# 5190-4641) per manufacturer’s instructions. Sequencing was performed on an Illumina HiSeq 2500 (Illumina cat.# SY–401–2501). DNA was sequenced to an average depth of 58. Quality control was performed with FastQC [51].

### Sequence alignment and processing

Adapter sequences were removed using CutAdapt, version 1.10 [52]. Low quality reads were trimmed using Sickle version 1.33 [53], in paired end mode with quality threshold set to 20 and length threshold set to 50 base pairs. Reads were aligned to the mm10 reference genome using the Burrows-Wheeler MEM aligner (BWA-MEM), version 0.7.12 [54]. Following alignment, duplicate reads were marked using Picard (https://broadinstitute.github.io/picard/) version 2.0.1. Base quality scores were then recalibrated using GATK version 3.6.0 [55–57]. Lists of known single nucleotide polymorphisms and indels for the FVB/N mouse strain, were downloaded from the Mouse Genomes Project, dbSNP release 142 and used for base quality score recalibration and mutation filtering (available at *ftp://ftp-mouse.sanger.ac.uk/*) [58].

### Mutation calling and analysis

Somatic mutations were called using the MuTect2 algorithm in GATK [55–57] with a minimum allowed base quality score of 20. Mutations were filtered against variants found in matched normal liver tissue and known single nucleotide polymorphisms for the FVB/N mouse strain. Candidate somatic mutations which did not pass the standard MuTect2 filters were removed from further analysis. Mutations not meeting the following requirements were also removed from further analysis: minimum allele depth of 10, minimum allele frequency of 0.05, and presence of the mutation in both forward and reverse strands. Mutations were annotated using SnpEff [59] and filtered for downstream analysis using SnpSift [60]. Candidate driver mutations were defined as moderate or high impact mutations, as defined by SnpEff, in driver genes as identified by the COSMIC cancer gene census [20]. To identify hotspot mutations, mouse amino acid positions were aligned to the orthologous human amino acid position using Clustal Omega [61] through UniProtKB [62] and used to query mutations found in the COSMIC database [20]. Mutational spectrum and signature analysis was performed using the deconstructSigs framework [63] modified to allow the use of the mm10 mouse reference genome. The COSMIC mutational signatures were used for reference [23].

### Copy number aberration analyses

Copy number aberrations were identified from exome sequence data using EXCAVATOR2 [64] using the mm10 reference genome. CNA calling was performed using standard settings, and a window size of 20000 base pairs. To identify potential driver CNAs, we filtered for CNAs associated with cancer in the COSMIC gene list [20].

### Analyses of human breast cancer data

Processed data from the METABRIC [6,7] and TCGA [22] cohorts were downloaded from, or analyzed directly on the cBioportal platform [65,66].

### Plot generation

Plots were created using R version 3.3.2 [46]. Heatmaps were created using ComplexHeatmap [67]. Mutational spectra histograms were created using the deconstructSigs package [63]. All other plots were generated using the ggplot2 package [68].

## Acknowledgements

We thank Phuong Vu, Eldri Undlien Due and Tina Brinks for helping with laboratory work, Prof. Rune Toftgård for providing the transgenic mouse lines, and the support staff at the Department of Comparative Medicine, Oslo University Hospital Radiumhospitalet for help with animal work. We are grateful to the members of the Department of Cancer Genetics, Institute for Cancer Research, Oslo University Hospital for insightful discussions, and in particular thank Tonje G. Lien for statistical input.

## Supporting information

**S1 Fig. The mutational spectra and mutational signatures of MPA/DMBA-induced mammary tumors A** T>A transversions were the most frequent mutation type in MPA/DMBA-induced tumors, followed by C>A transversions. **B** Heatmap of mutational frequencies by trinucleotide context. There was an overrepresentation of T>N mutations in positions with a 3’ guanine and C>N mutations in positions with a 3’ adenine. **C** Histogram of C>A and T>A transversions by trinucleotide context in a representative tumor (*S159_14_8*). **D** Mutation signature 22 was the predominant mutational signature in the MPA/DMBA-induced tumors and was evident in all tumors in the cohort.

**S2 Fig. Gene expression scores by cluster in MPA/DMBA-induced tumors** Expression scores by cluster for genes related to differentiation, adhesion, luminal features, proliferation, vascular content, EMT, immune features, interferon signaling and immunosuppression. Two-tailed Wilcoxon rank-sum test. ns = not significant, p > 0.05. * = p < 0.05. ** = p < 0.01. *** = p < 0.001.

**S3 Fig. Expression of *Cd24a* and *Cd44* by cluster in MPA/DMBA-induced tumors** Claudin-low-like tumors had a lower expression of *Cd24a* and a higher expression of *Cd44* compared to the mixed cluster of tumors (p = 0.003 and p = 0.005, respectively, two-tailed, Wilcoxon rank-sum test), indicating a stem cell-like phenotype in the claudin-low-like tumors.

S1 File. Mouse characteristics and histopathological data

S2 File. Subtype correlations for MPA/DMBA-induced tumors

S3 File. Mutations observed in MPA/DMBA-induced tumors

S4 File. MPA/DMBA-induced tumor driver gene mutations in the COSMIC database

S5 File. Comparative mutation rates in MPA/DMBA-induced tumors and human breast tumors in the TCGA cohort

S6 File. Mutational signatures for all MPA/DMBA-induced tumors

S7 File. Gene lists used for gene expression scores

## References

1. Herschkowitz JI, Simin K, Weigman VJ, Mikaelian I, Usary J, Hu Z, et al. Identification of conserved gene expression features between murine mammary carcinoma models and human breast tumors. Genome Biol. 2007;8(5):R76.

2. Prat A, Parker JS, Karginova O, Fan C, Livasy C, Herschkowitz JI, et al. Phenotypic and molecular characterization of the claudin-low intrinsic subtype of breast cancer. Breast Cancer Res. 2010;12(5):R68.

3. Sabatier R, Finetti P, Guille A, Adelaide J, Chaffanet M, Viens P, et al. Claudin-low breast cancers: clinical, pathological, molecular and prognostic characterization. Mol Cancer. 2014;13(1):228.

4. Prat A, Perou CM. Deconstructing the molecular portraits of breast cancer. Mol Oncol. 2011;5(1):5–23.

5. Dias K, Dvorkin-Gheva A, Hallett RM, Wu Y, Hassell J, Pond GR, et al. Claudin-low breast cancer; clinical & pathological characteristics. PLoS One. 2017;12(1):e0168669.

6. Curtis C, Shah SP, Chin S-F, Turashvili G, Rueda OM, Dunning MJ, et al. The genomic and transcriptomic architecture of 2,000 breast tumours reveals novel subgroups. Nature. 2012;486(7403):346.

7. Pereira B, Chin S-F, Rueda OM, Vollan H-KM, Provenzano E, Bardwell HA, et al. The somatic mutation profiles of 2,433 breast cancers refine their genomic and transcriptomic landscapes. Nat Commun. 2016;7:11479.

8. Hennessy BT, Gonzalez-Angulo A-M, Stemke-Hale K, Gilcrease MZ, Krishnamurthy S, Lee J-S, et al. Characterization of a naturally occurring breast cancer subset enriched in epithelial-to-mesenchymal transition and stem cell characteristics. Cancer Res. 2009;69(10):4116–24.

9. Prat A, Adamo B, Cheang MCU, Anders CK, Carey LA, Perou CM. Molecular characterization of basal-like and non-basal-like triple-negative breast cancer. Oncologist. 2013;18(2):123–33.

10. Aldaz CM, Liao QY, LaBate M, Johnston DA. Medroxyprogesterone acetate accelerates the development and increases the incidence of mouse mammary tumors induced by dimethylbenzanthracene. Carcinogenesis. 1996;17(9):2069–72.

11. Pfefferle AD, Herschkowitz JI, Usary J, Harrell J, Spike BT, Adams JR, et al. Transcriptomic classification of genetically engineered mouse models of breast cancer identifies human subtype counterparts. Genome Biol. 2013;14(11):R125.

12. Yin Y, Bai R, Russell RG, Beildeck ME, Xie Z, Kopelovich L, et al. Characterization of medroxyprogesterone and DMBA-induced multilineage mammary tumors by gene expression profiling. Mol Carcinog Publ Coop with Univ Texas MD Anderson Cancer Cent. 2005;44(1):42–50.

13. Gonzalez-Suarez E, Jacob AP, Jones J, Miller R, Roudier-Meyer MP, Erwert R, et al. RANK ligand mediates progestin-induced mammary epithelial proliferation and carcinogenesis. Nature. 2010;468(7320):103.

14. Baird WM, Hooven LA, Mahadevan B. Carcinogenic polycyclic aromatic hydrocarbon-DNA adducts and mechanism of action. Environ Mol Mutagen. 2005;45(2–3):106–14.

15. Frenkel K. 7,12-dimethylbenz[a]anthracene induces oxidative DNA modification in vivo. Free Radic Biol Med. 1995;19(3):373–80.

16. Dean JH, Ward EC, Murray MJ, Lauer LD, House R V. Mechanisms of dimethylbenzanthracene-induced immunotoxicity. Clin Physiol Biochem. 1985;3(2– 3):98–110.

17. Miyata M, Furukawa M, Takahashi K, Gonzalez FJ, Yamazoe Y. Mechanism of 7, 12-dimethylbenz[a]anthracene-induced immunotoxicity: role of metabolic activation at the target organ. Jpn J Pharmacol. 2001;86(3):302–9.

18. Trichopoulos D, Adami H, Ekbom A, Hsieh C, Lagiou P. Early life events and conditions and breast cancer risk: from epidemiology to etiology. Int J cancer. 2008;122(3):481–5.

19. Lawrence MS, Stojanov P, Polak P, Kryukov G V, Cibulskis K, Sivachenko A, et al. Mutational heterogeneity in cancer and the search for new cancer-associated genes. Nature. 2013;499(7457):214–8.

20. Forbes SA, Beare D, Boutselakis H, Bamford S, Bindal N, Tate J, et al. COSMIC: somatic cancer genetics at high-resolution. Nucleic Acids Res. 2016;45(D1):D777–83.

21. Abba MC, Zhong Y, Lee J, Kil H, Lu Y, Takata Y, et al. DMBA induced mouse mammary tumors display high incidence of activating Pik3caH1047 and loss of function Pten mutations. Oncotarget. 2016;5.

22. The Cancer Genome Atlas Network. Comprehensive molecular portraits of human breast tumours. Nature. 2012;490(7418):61.

23. Alexandrov LB, Nik-Zainal S, Wedge DC, Aparicio S a. JR, Behjati S, Biankin A V, et al. Signatures of mutational processes in human cancer. Nature. 2013;500(7463):415–21.

24. Ciriello G, Miller ML, Aksoy BA, Senbabaoglu Y, Schultz N, Sander C. Emerging landscape of oncogenic signatures across human cancers. Nat Genet. 2013;45(10):1127–33.

25. Prat A, Parker JS, Karginova O, Fan C, Livasy C, Herschkowitz JI, et al. Phenotypic and molecular characterization of the claudin-low intrinsic subtype of breast cancer. Breast Cancer Res. 2010;12(5):R68.

26. Visvader JE, Stingl J. Mammary stem cells and the differentiation hierarchy: current status and perspectives. Genes Dev. 2014;28(11):1143–58.

27. Hegde PS, Karanikas V, Evers S. The Where, the When, and the How of Immune Monitoring for Cancer Immunotherapies in the Era of Checkpoint Inhibition. Clin Cancer Res. 2016 Apr;22(8):1865–74.

28. Jamieson NB, Maker A V. Gene-expression profiling to predict responsiveness to immunotherapy. Nat Publ Gr. 2016;24(3):134–40.

29. Zelenay S, Van Der Veen AG, Böttcher JP, Snelgrove KJ, Rogers N, Acton SE, et al. Cyclooxygenase-dependent tumor growth through evasion of immunity. Cell. 2015;162(6):1257–70.

30. Chokr N, Chokr S. Immune Checkpoint Inhibitors in Triple Negative Breast Cancer: What is the Evidence? 2018;

31. Prat A, Perou CM. Mammary development meets cancer genomics. Nat Med. 2009;15(8):842.

32. Hanahan D, Coussens LM. Accessories to the crime: functions of cells recruited to the tumor microenvironment. Cancer Cell. 2012;21(3):309–22.

33. Hollern DP, Swiatnicki MR, Andrechek ER. Histological subtypes of mouse mammary tumors reveal conserved relationships to human cancers. PLoS Genet. 2018;14(1):e1007135.

34. Rennhack J, Swiatnicki M, Zhang Y, Li C, Bylett E, Ross C, et al. Integrated sequence and gene expression analysis of mouse models of breast cancer reveals critical events with human parallels. bioRxiv. 2018; 375154.

35. Morel A-P, Ginestier C, Pommier RM, Cabaud O, Ruiz E, Wicinski J, et al. A stemness-related ZEB1–MSRB3 axis governs cellular pliancy and breast cancer genome stability. Nat Med. 2017;23(5):568.

36. Burstein MD, Tsimelzon A, Poage GM, Covington KR, Contreras A, Fuqua SAW, et al. Comprehensive genomic analysis identifies novel subtypes and targets of triple-negative breast cancer. Clin Cancer Res. 2015;21(7):1688–98.

37. Weigelt B, Ng CKY, Shen R, Popova T, Schizas M, Natrajan R, et al. Metastatic breast carcinomas display genomic and transcriptomic heterogeneity. Mod Pathol. 2015;28(3):340.

38. Taylor NA, Vick SC, Iglesia MD, Brickey WJ, Midkiff BR, McKinnon KP, et al. Treg depletion potentiates checkpoint inhibition in claudin-low breast cancer. J Clin Invest. 2017 Aug;127(9):3472–83.

39. Zarghi A, Arfaei S. Selective COX-2 Inhibitors: A Review of Their Structure-Activity Relationships. Iran J Pharm Res IJPR. 2011;10(4):655–83.

40. Dannenberg AJ, DuBois RN. COX-2?: a new target for cancer prevention and treatment. Karger; 2003. 291 p.

41. Tsujii M, DuBois RN. Alterations in cellular adhesion and apoptosis in epithelial cells overexpressing prostaglandin endoperoxide synthase 2. Cell. 1995 Nov;83(3):493–501.

42. Basu GD, Liang WS, Stephan DA, Wegener LT, Conley CR, Pockaj BA, et al. A novel role for cyclooxygenase-2 in regulating vascular channel formation by human breast cancer cells. Breast Cancer Res. 2006;8(6):R69.

43. Chuck Harrell J, Pfefferle AD, Zalles N, Prat A, Fan C, Khramtsov A, et al. Endothelial-like properties of claudin-low breast cancer cells promote tumor vascular permeability and metastasis. Clin Exp Metastasis. 2014;31:33–45.

44. Yan X, Zhang S, Deng Y, Wang P, Hou Q, Xu H. Prognostic Factors for Checkpoint Inhibitor Based Immunotherapy: An Update With New Evidences. Front Pharmacol. 2018 Sep;9:1050.

45. Norum JH, Bergström Å, Andersson AB, Kuiper R V, Hoelzl MA, Sørlie T, et al. A conditional transgenic mouse line for targeted expression of the stem cell marker LGR5. Dev Biol. 2015;404(2):35–48.

46. Team RC, Computing RF for S. R: A language and environment for statistical computing. Vienna, Austria: R Foundation for Statistical Computing; 2017. p.

47. Yoshihara K, Shahmoradgoli M, Martínez E, Vegesna R, Kim H, Torres-Garcia W, et al. Inferring tumour purity and stromal and immune cell admixture from expression data. Nat Commun. 2013;4:2612.

48. Kardos J, Chai S, Mose LE, Selitsky SR, Krishnan B, Saito R, et al. Claudin-low bladder tumors are immune infiltrated and actively immune suppressed. JCI insight. 2016;1(3):e85902.

49. Nielsen TO, Parker JS, Leung S, Voduc D, Ebbert M, Vickery T, et al. A Comparison of PAM50 Intrinsic Subtyping with Immunohistochemistry and Clinical Prognostic Factors in Tamoxifen-Treated Estrogen Receptor-Positive Breast Cancer. Clin Cancer Res. 2010 Nov;16(21):5222–32.

50. Subramanian A, Tamayo P, Mootha VK, Mukherjee S, Ebert BL, Gillette MA, et al. Gene set enrichment analysis: a knowledge-based approach for interpreting genome-wide expression profiles. Proc Natl Acad Sci. 2005;102(43):15545–50.

51. Andrews S. FastQC: A quality control tool for high throughput sequence data. Ref Source. 2010;

52. Martin M. Cutadapt removes adapter sequences from high-throughput sequencing reads. EMBNet.journal. 2011;17(1).

53. Joshi NA FJN. Sickle: A sliding-window, adaptive, quality-based tool for FastQ files. 2011;

54. Li H. Aligning sequence reads, clone sequences and assembly contigs with BWA-MEM. arXiv:13033997. 2013;

55. Van der Auwera GA, Carneiro MO, Hartl C, Poplin R, del Angel G, Levy-Moonshine A, et al. From FastQ Data to High-Confidence Variant Calls: The Genome Analysis Toolkit Best Practices Pipeline. In: Current protocols in bioinformatics. John Wiley & Sons, Inc.; 2013.

56. McKenna A, Hanna M, Banks E, Sivachenko A, Cibulskis K, Kernytsky A, et al. The Genome Analysis Toolkit: a MapReduce framework for analyzing next-generation DNA sequencing data. Genome Res. 2010 Sep;20(9):1297–303.

57. DePristo MA, Banks E, Poplin R, Garimella K V, Maguire JR, Hartl C, et al. A framework for variation discovery and genotyping using next-generation DNA sequencing data. Nat Genet. 2011;43(5):491–8.

58. Wong K, Bumpstead S, Van Der Weyden L, Reinholdt LG, Wilming LG, Adams DJ, et al. Sequencing and characterization of the FVB/NJ mouse genome. Genome Biol. 2012;13(8):1–12.

59. Cingolani P, Platts A, Wang LL, Coon M, Nguyen T, Wang L, et al. A program for annotating and predicting the effects of single nucleotide polymorphisms, SnpEff. Fly (Austin). 2012;6(2):80–92.

60. Cingolani P, Patel VM, Coon M, Nguyen T, Land SJ, Ruden DM, et al. Using Drosophila melanogaster as a Model for Genotoxic Chemical Mutational Studies with a New Program, SnpSift. Front Genet. 2012;3:35.

61. Sievers F, Wilm A, Dineen D, Gibson TJ, Karplus K, Li W, et al. Fast, scalable generation of high-quality protein multiple sequence alignments using Clustal Omega. Mol Syst Biol. 2014;7(1):539.

62. The UniProt Consortium. UniProt: the universal protein knowledgebase. Nucleic Acids Res. 2017;45(D1):D158–69.

63. Rosenthal R, McGranahan N, Herrero J, Taylor BS, Swanton C. deconstructSigs: delineating mutational processes in single tumors distinguishes DNA repair deficiencies and patterns of carcinoma evolution. Genome Biol. 2016;17(1):1.

64. D’Aurizio R, Pippucci T, Tattini L, Giusti B, Pellegrini M, Magi A. Enhanced copy number variants detection from whole-exome sequencing data using EXCAVATOR2. Nucleic Acids Res. 2016;44(20):e154–e154.

65. Gao J, Aksoy BA, Dogrusoz U, Dresdner G, Gross B, Sumer SO, et al. Integrative analysis of complex cancer genomics and clinical profiles using the cBioPortal. Sci Signal. 2013;6(269):pl1.

66. Cerami E, Gao J, Dogrusoz U, Gross BE, Sumer SO, Aksoy BA, et al. The cBio cancer genomics portal: an open platform for exploring multidimensional cancer genomics data. AACR; 2012.

67. Gu Z, Eils R, Schlesner M. Complex heatmaps reveal patterns and correlations in multidimensional genomic data. Bioinformatics. 2016;32(18):2847–9.

68. Wickham H. ggplot2. New York, NY: Springer New York; 2009.

