## Supplementary figures and images for "Claudin-low-like mouse mammary tumors show distinct transcriptomic patterns uncoupled from genomic drivers"

### S1Figure

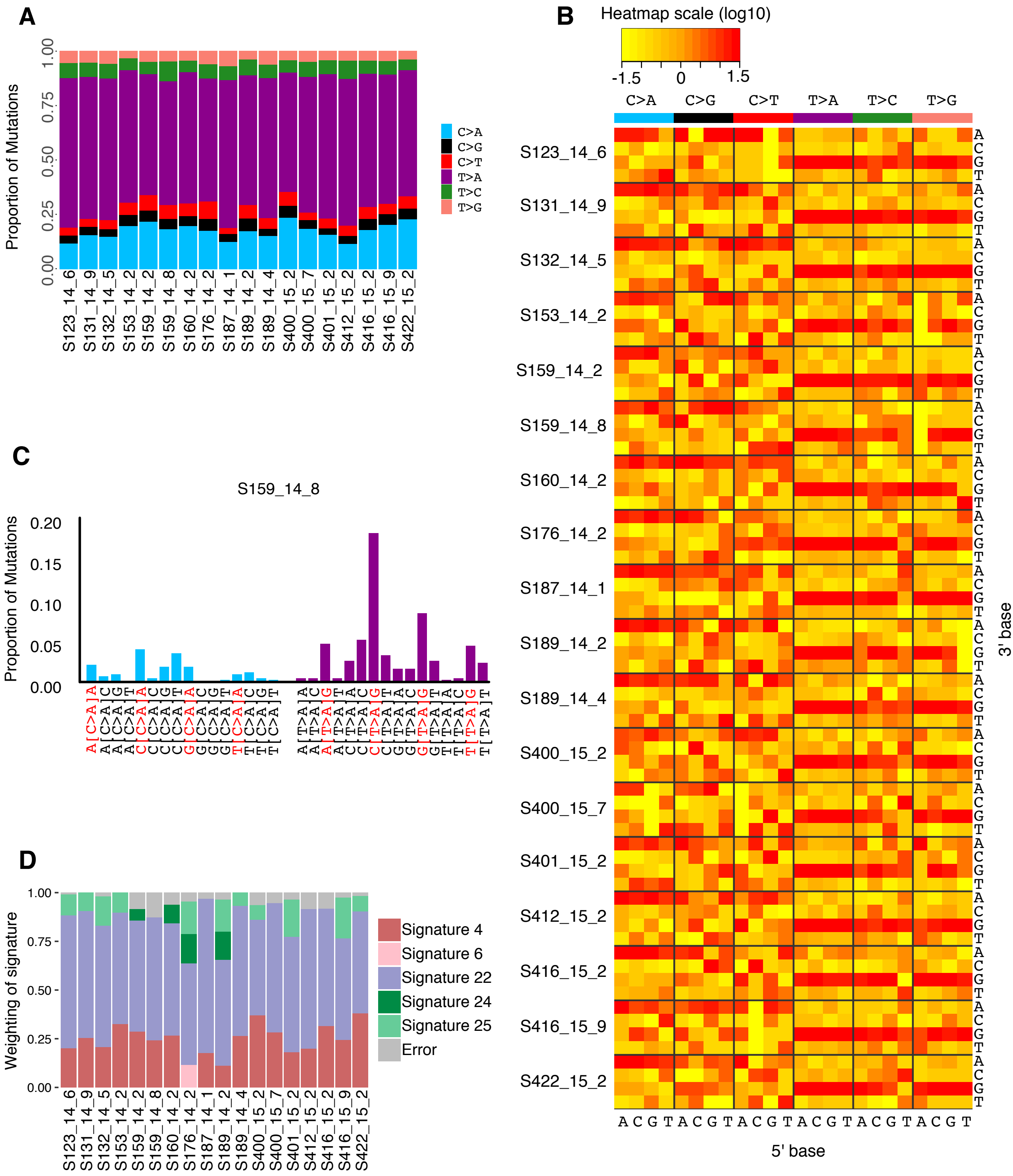

### S2Figure

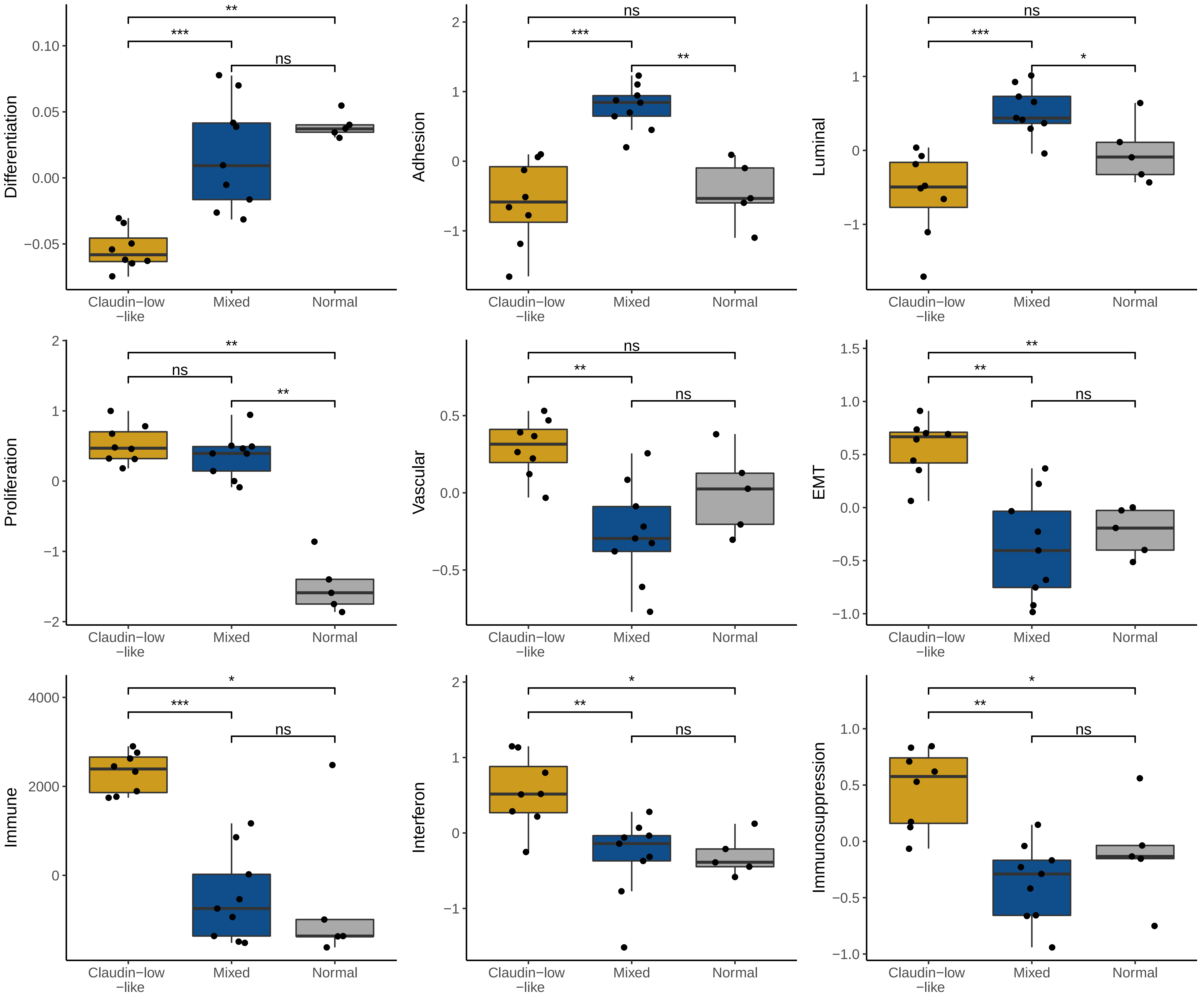

### S3Figure

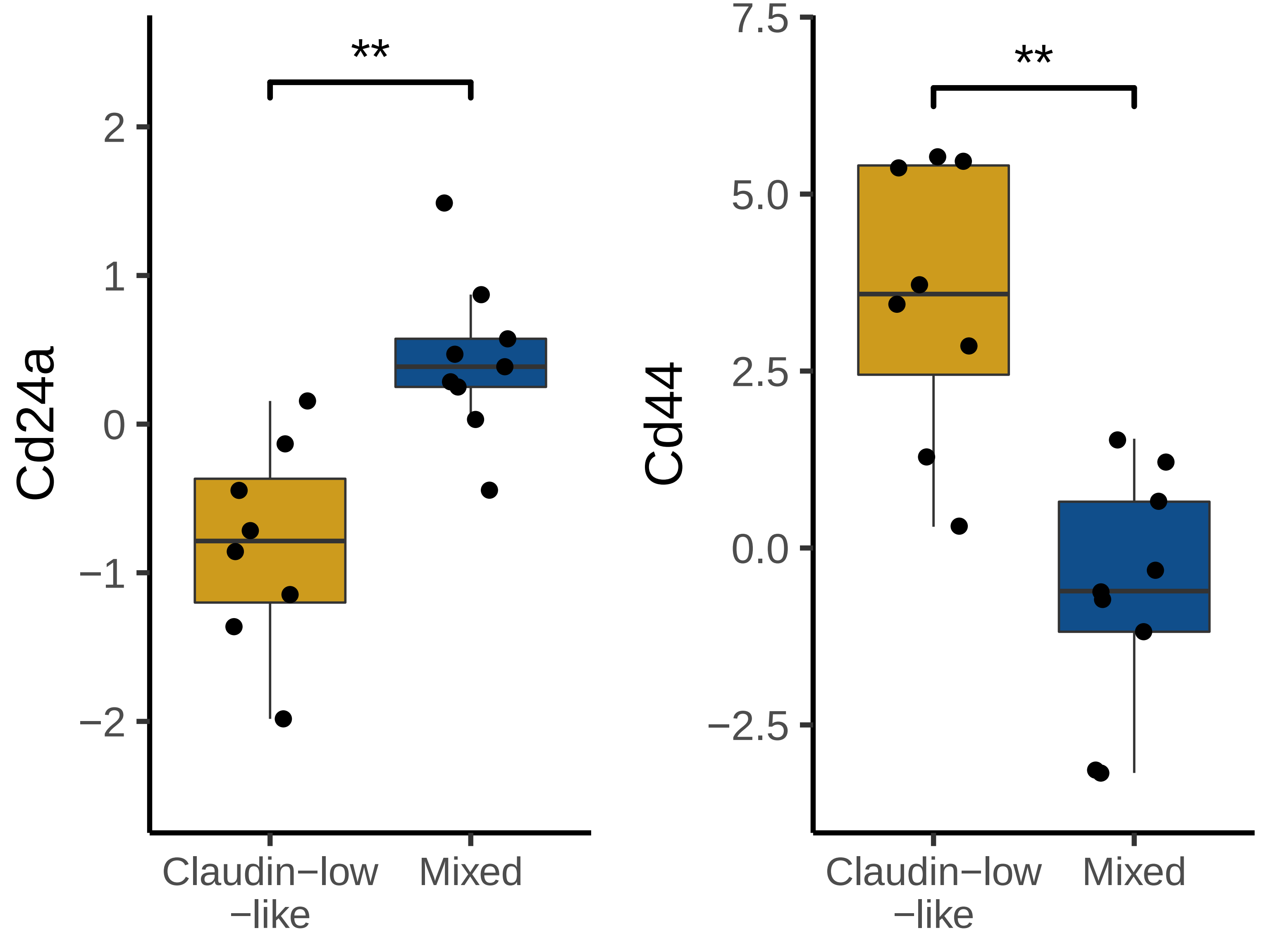

### S123_14_6.pdf

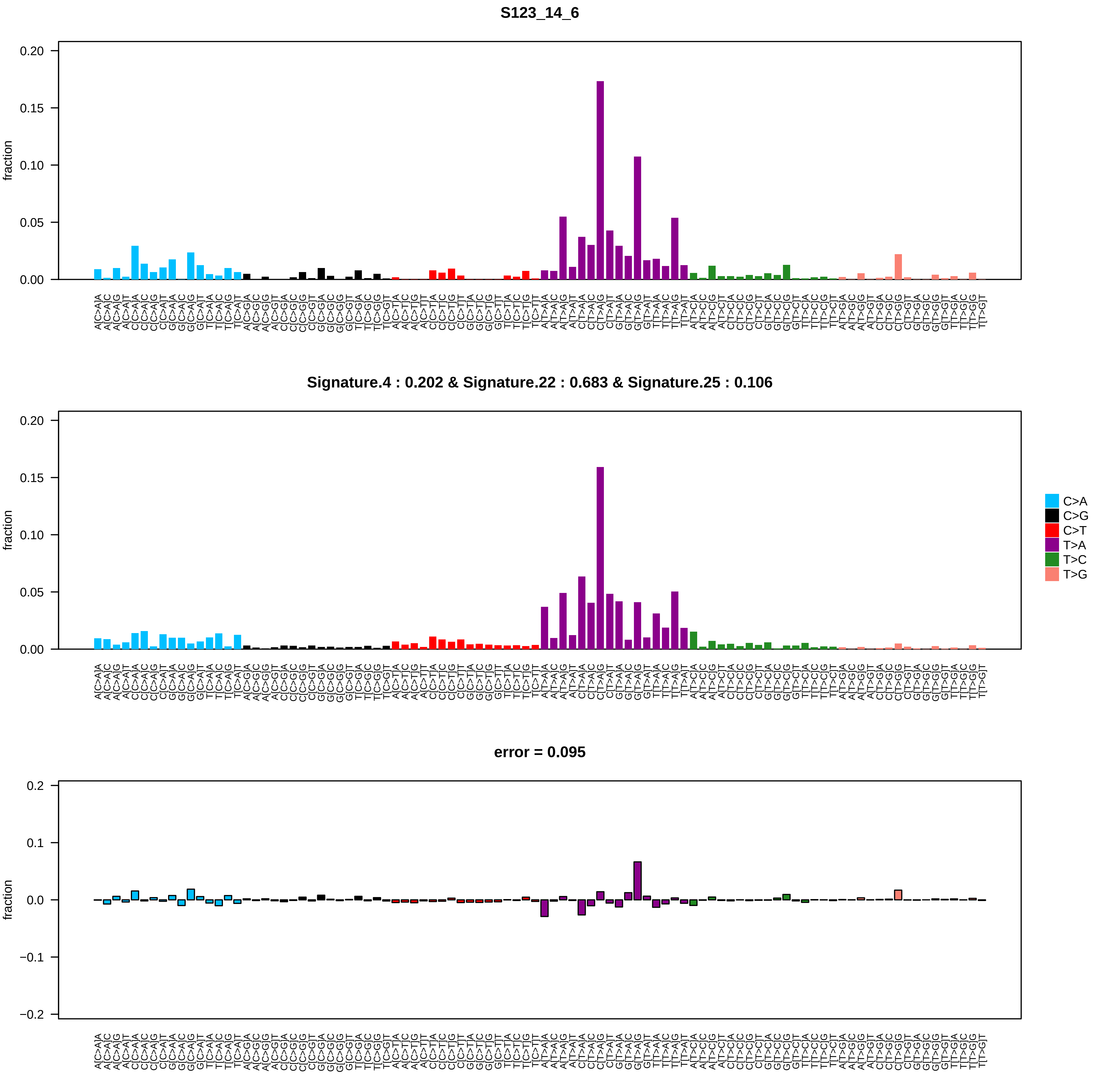

### S131_14_9.pdf

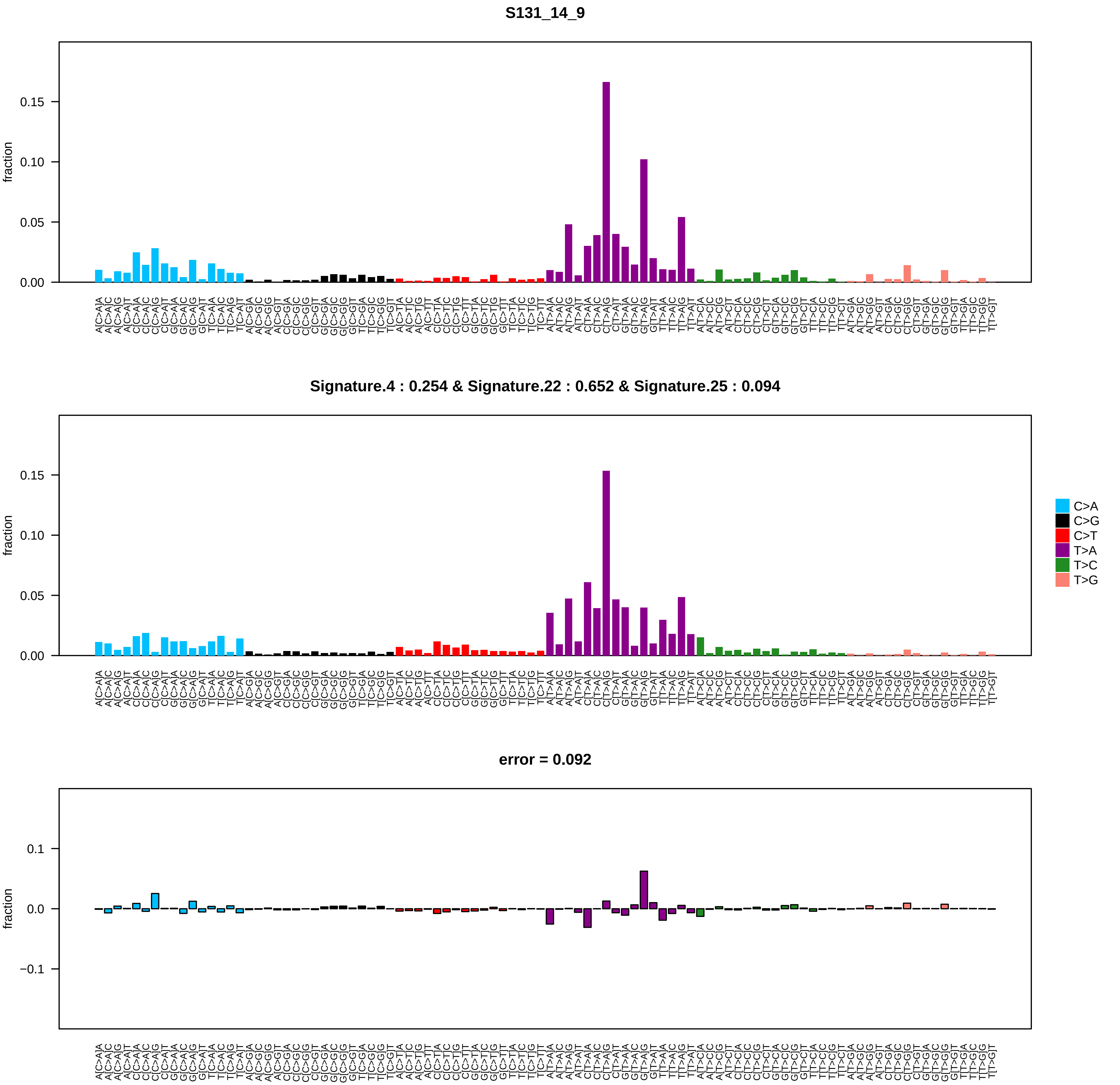

### S132_14_5.pdf

**S132_14_5**

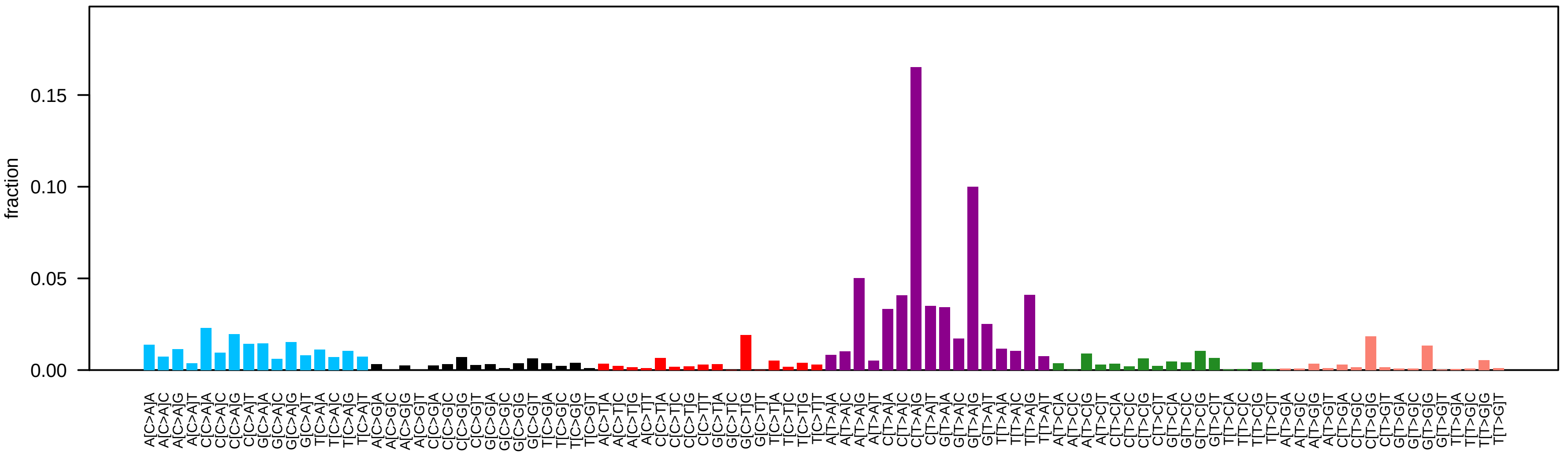


**Signature.4 : 0.209 & Signature.22 : 0.622 & Signature.25 : 0.151**

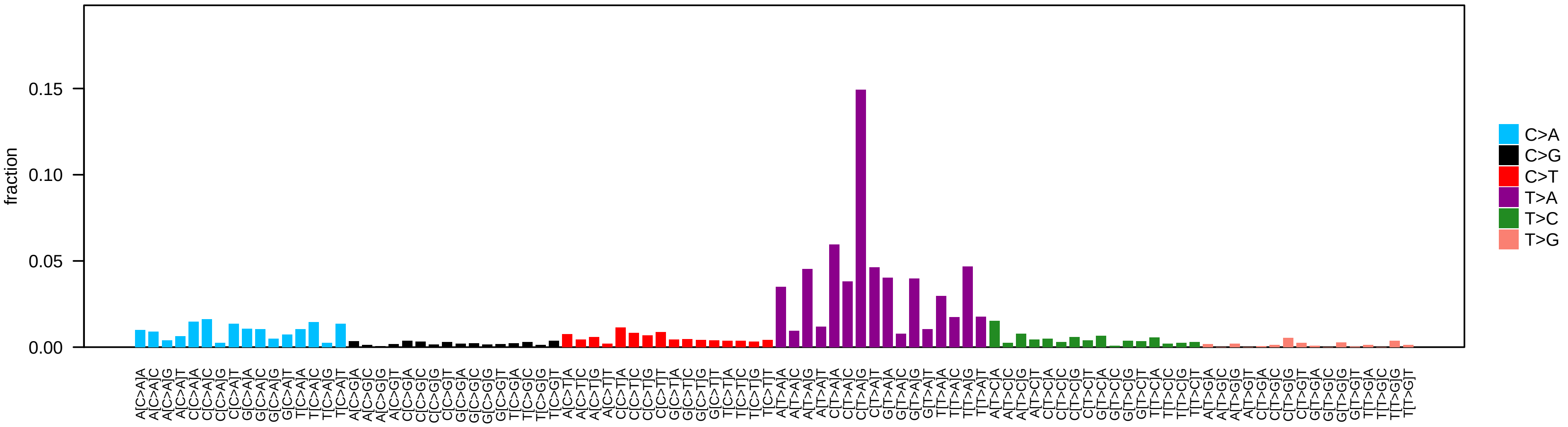


**error = 0.09**

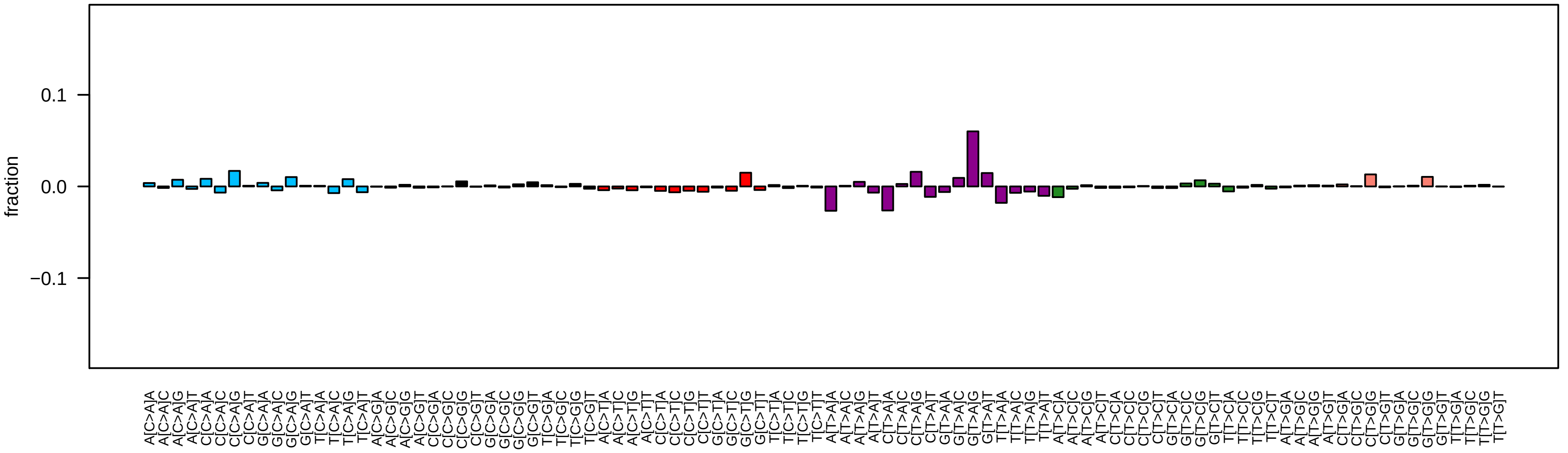

### S153_14_2.pdf

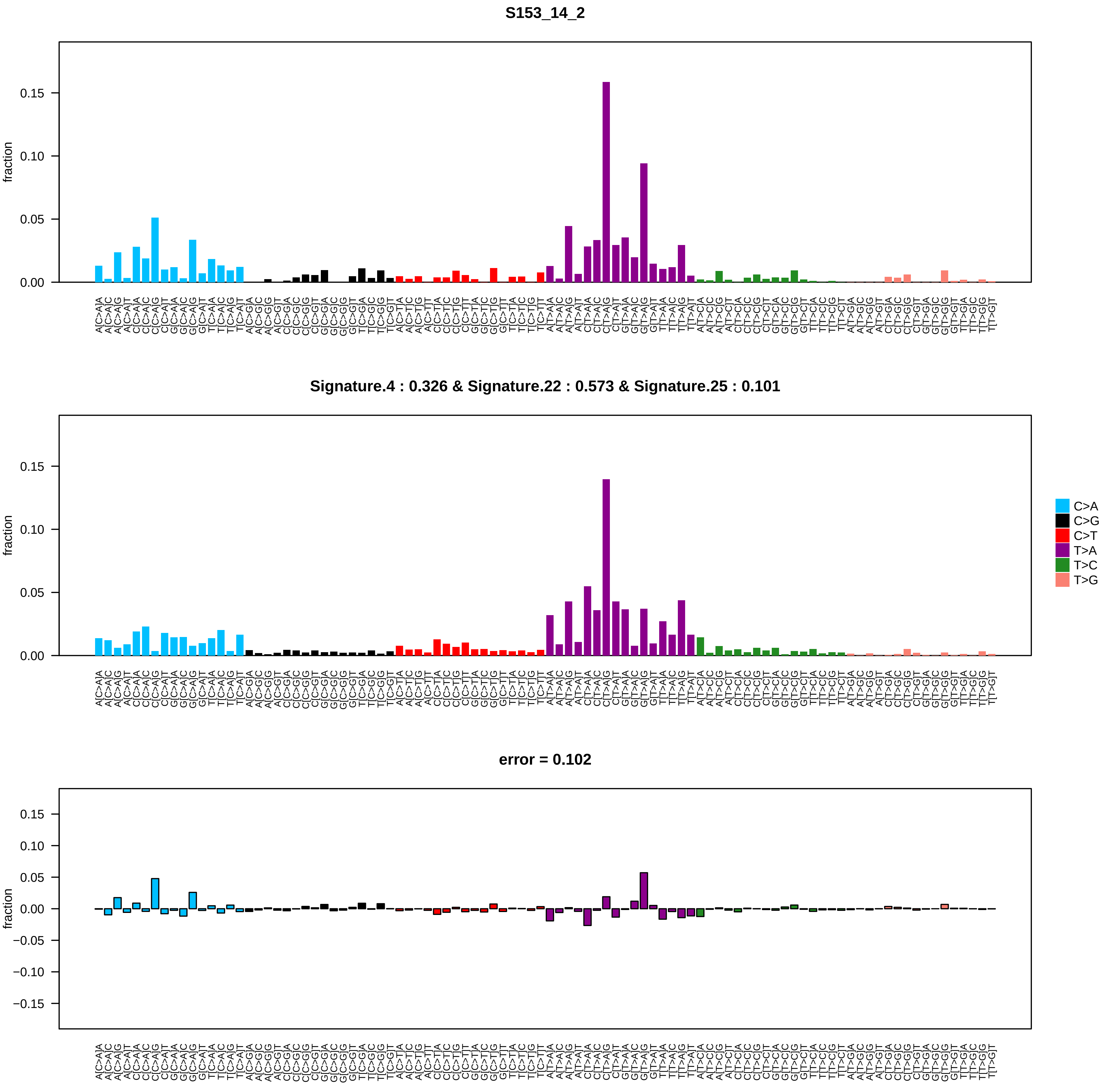

### S159_14_2.pdf

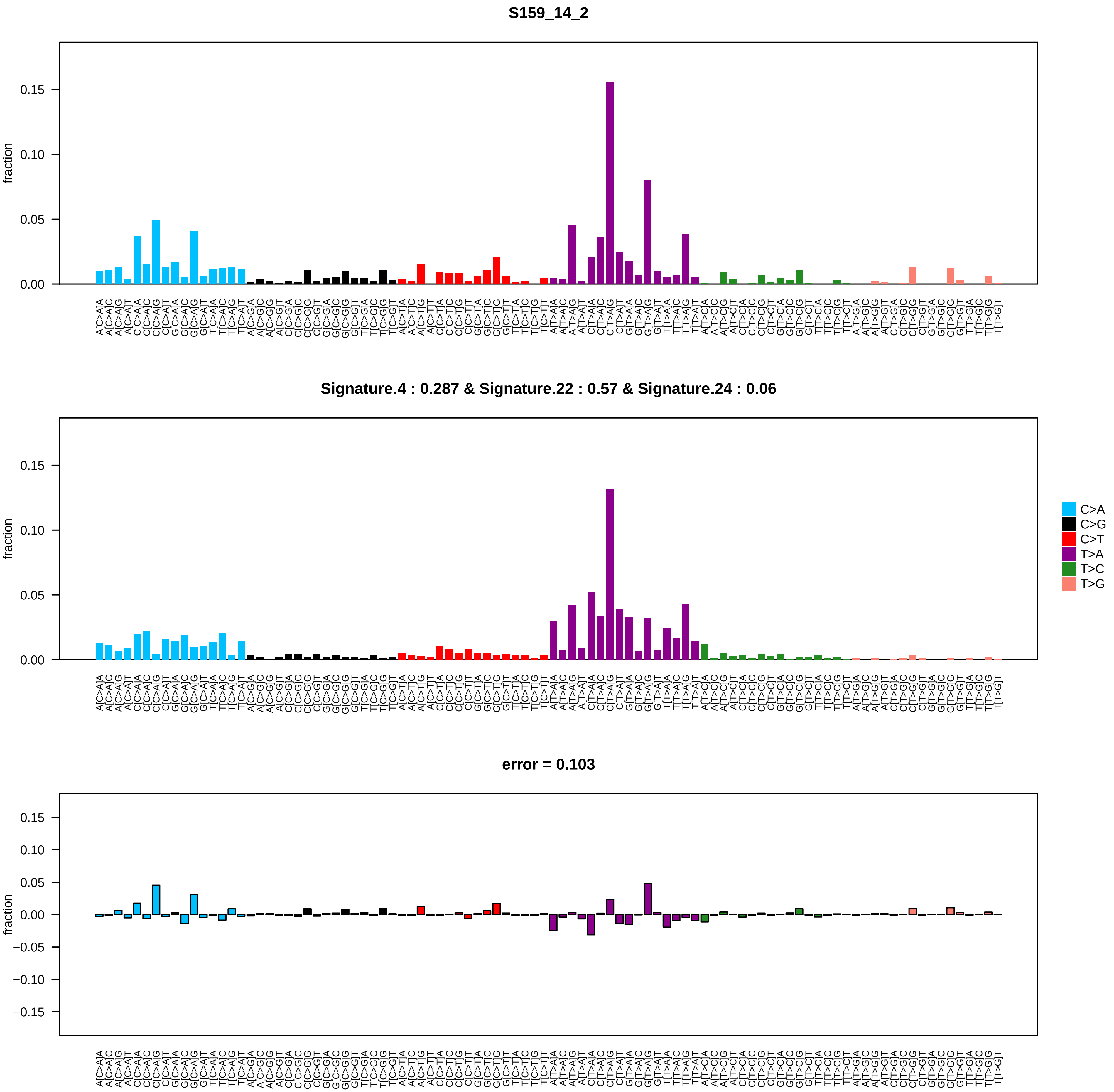

### S159_14_8.pdf

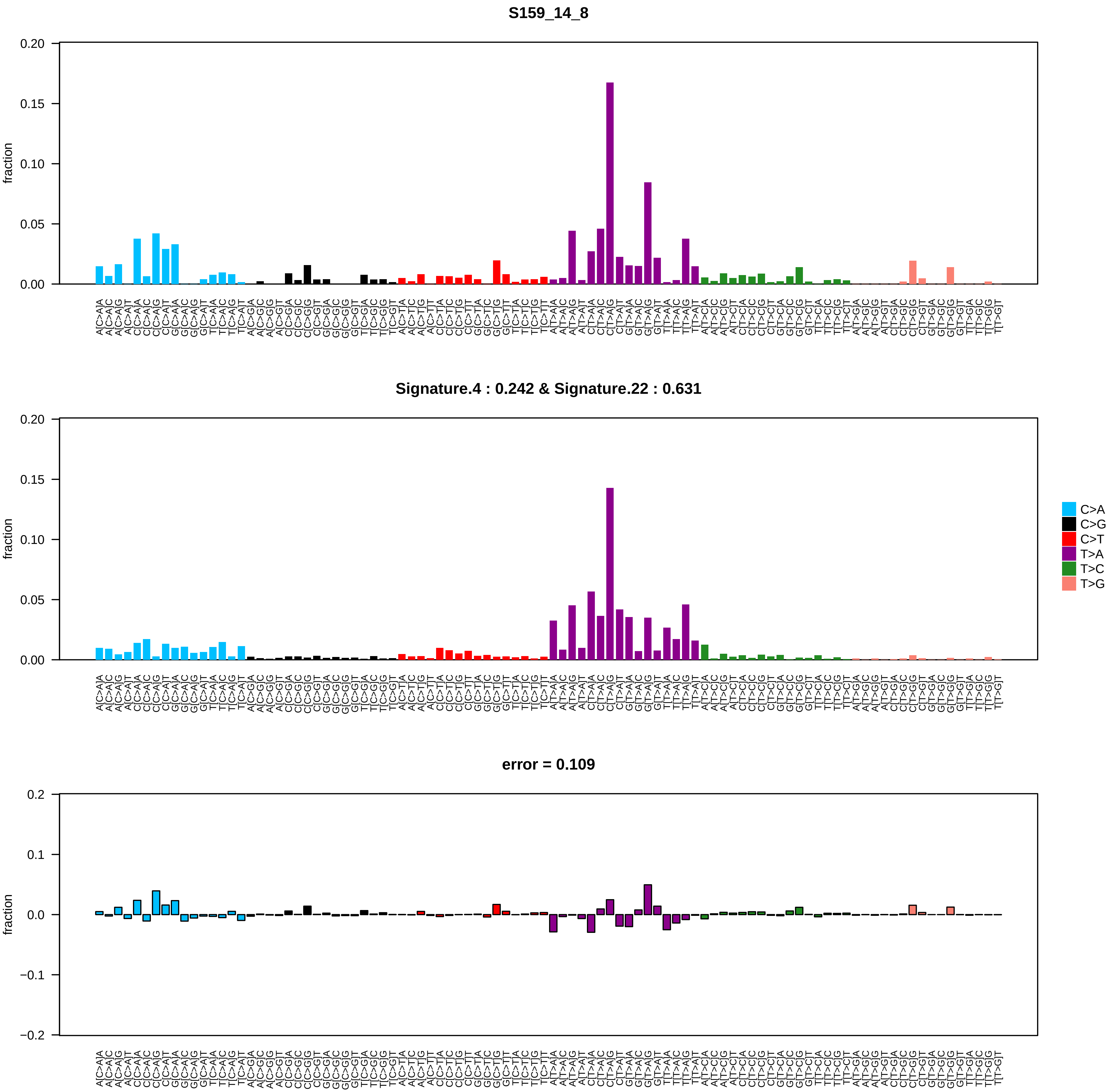

### S160_14_2.pdf

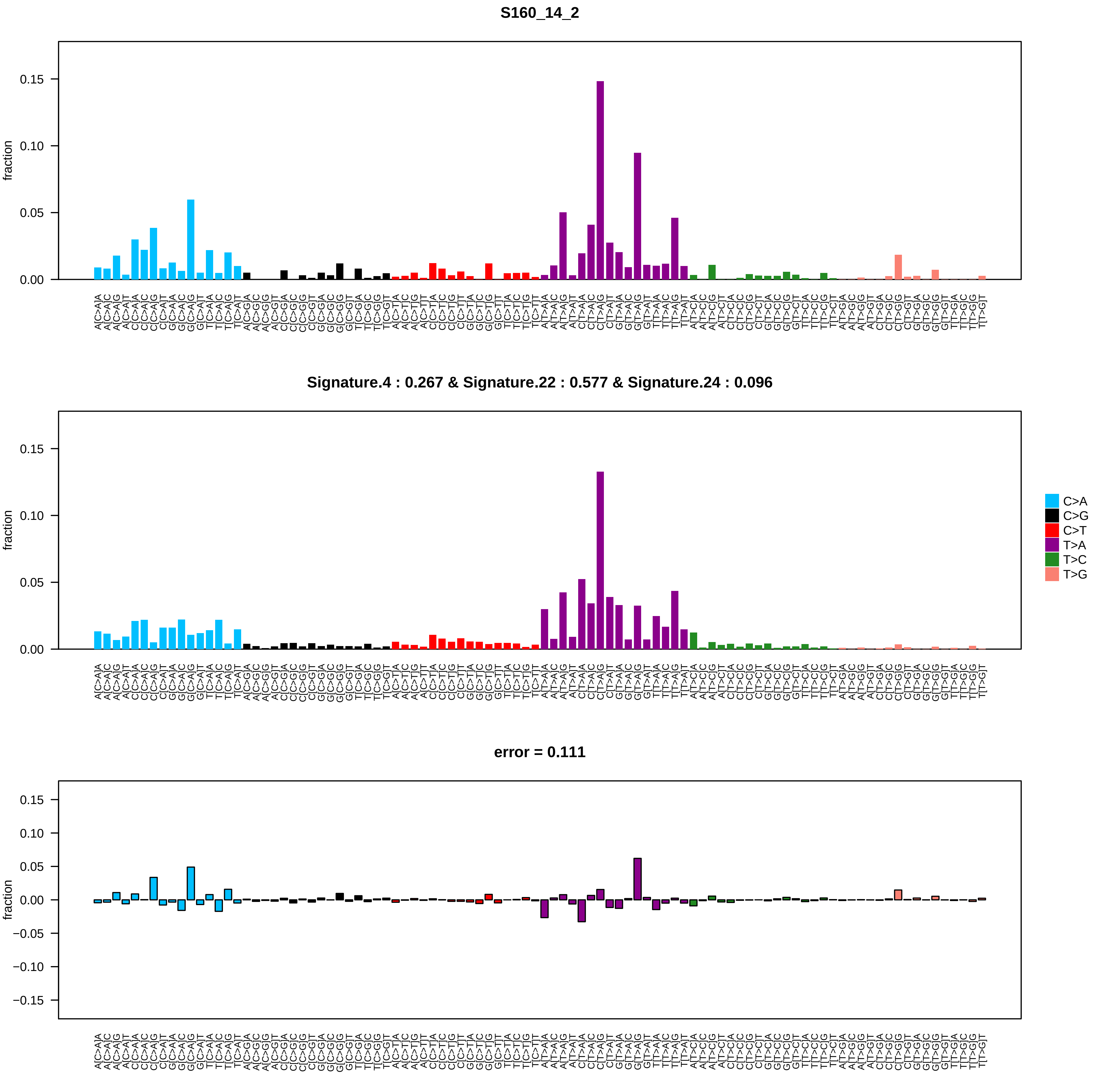

### S176_14_2.pdf

## S176_14_2

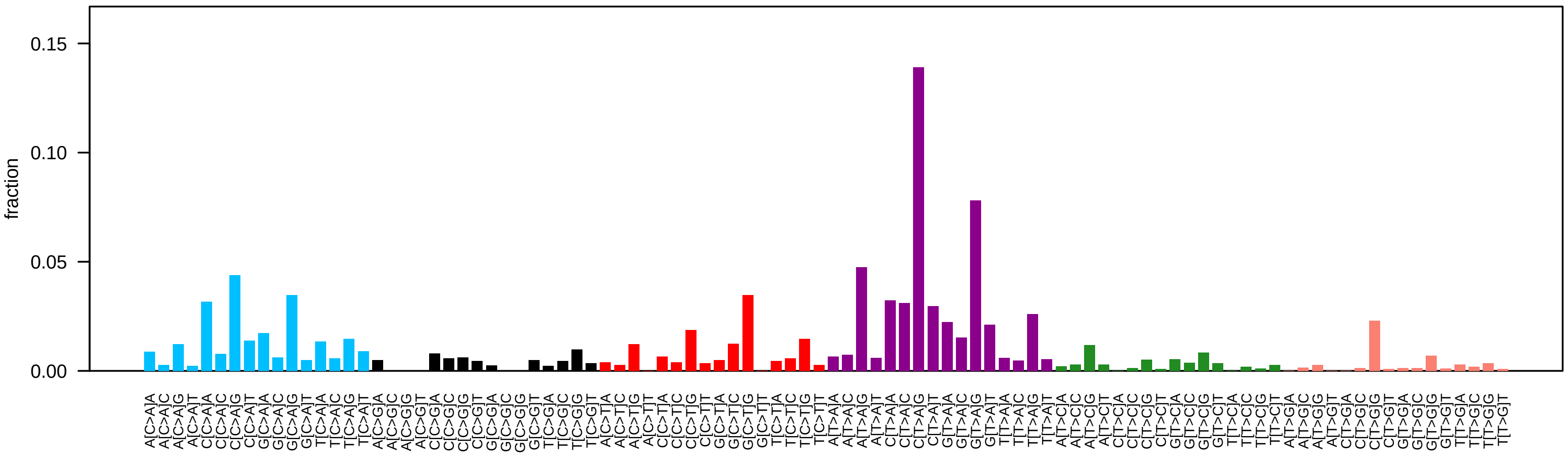


#### Signature.6 : 0.117 & Signature.22 : 0.52 & Signature.24 : 0.151 & Signature.25 : 0.168

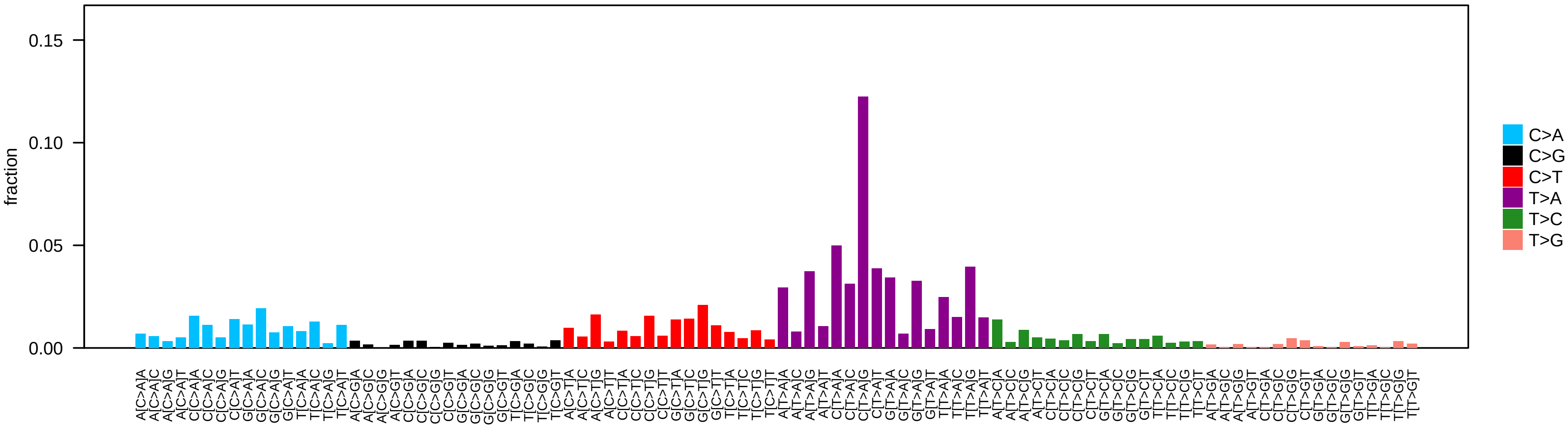


#### error = 0.094

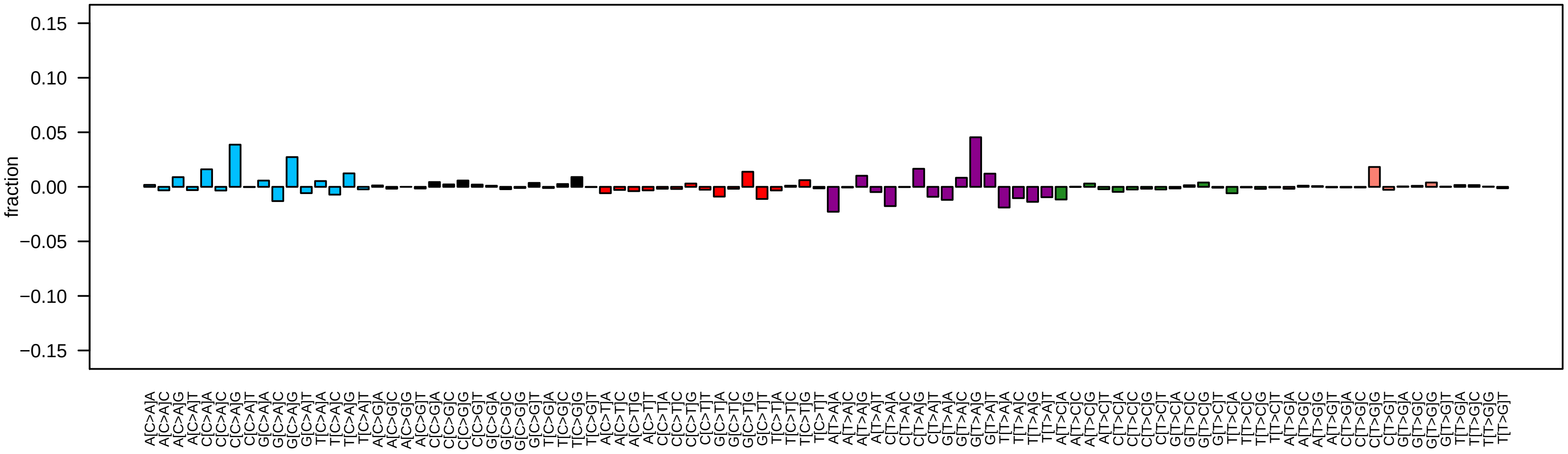

### S187_14_1.pdf

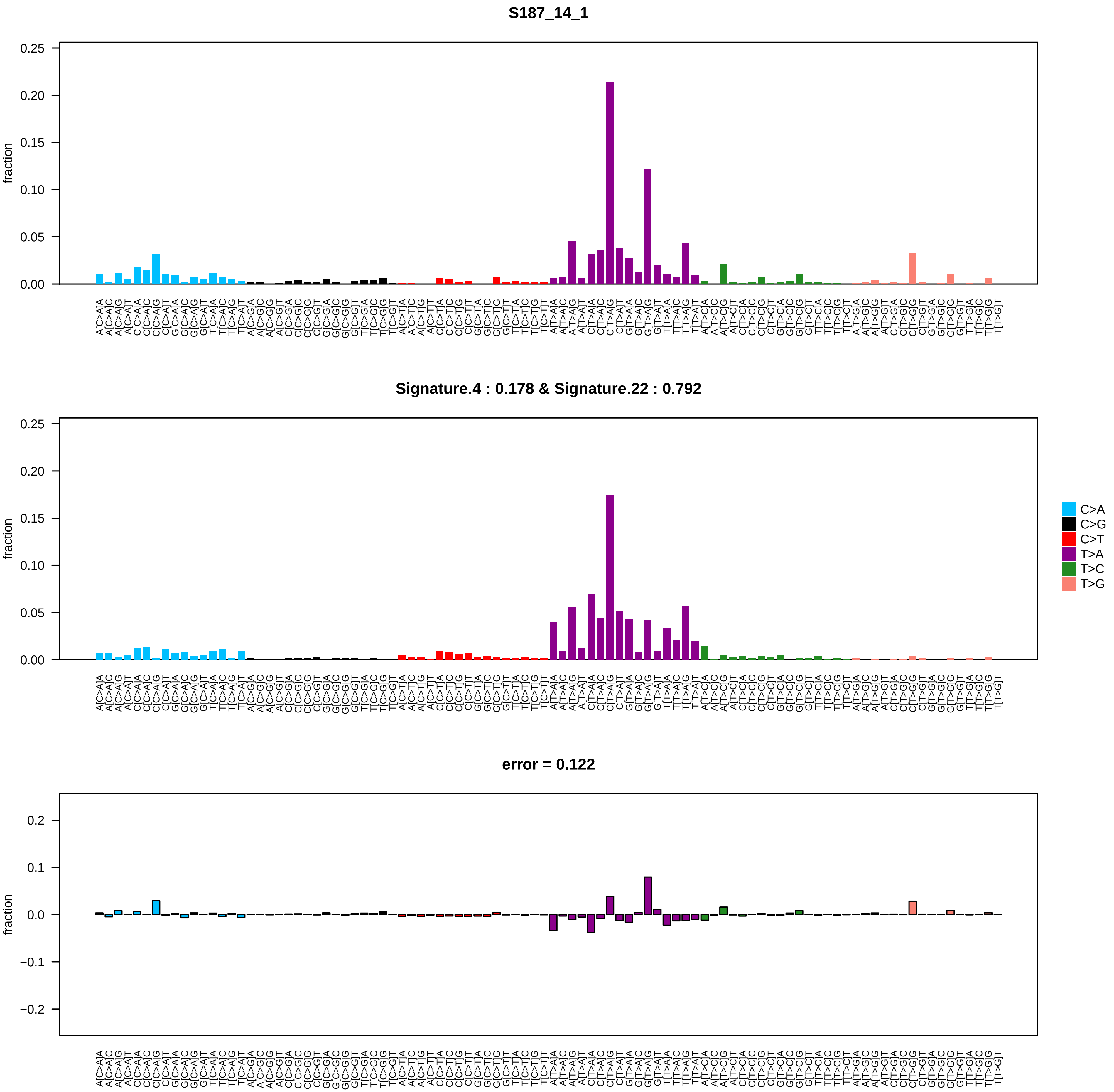

### S189_14_2.pdf

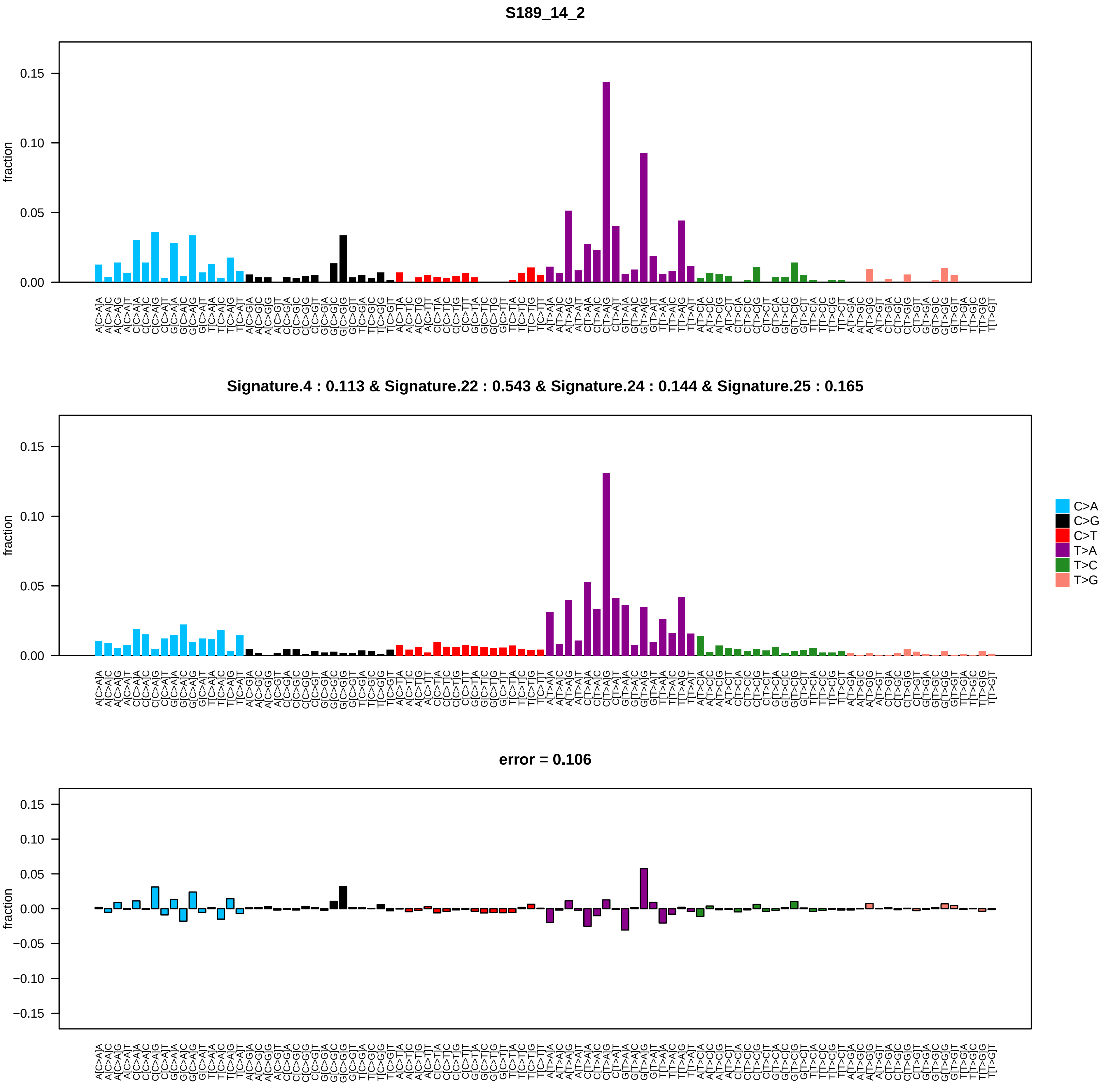

### S189_14_4.pdf

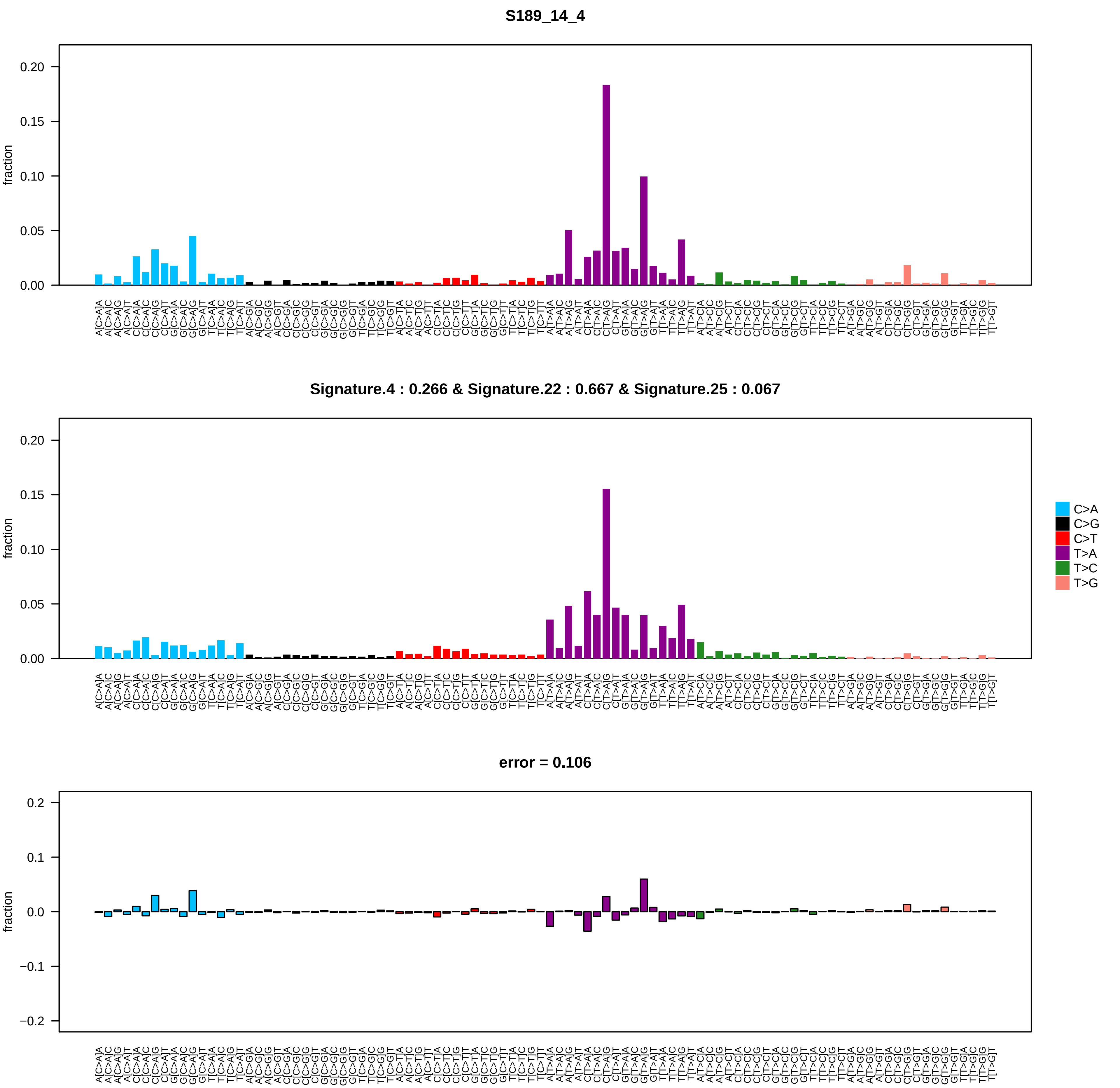

### S400_15_2.pdf

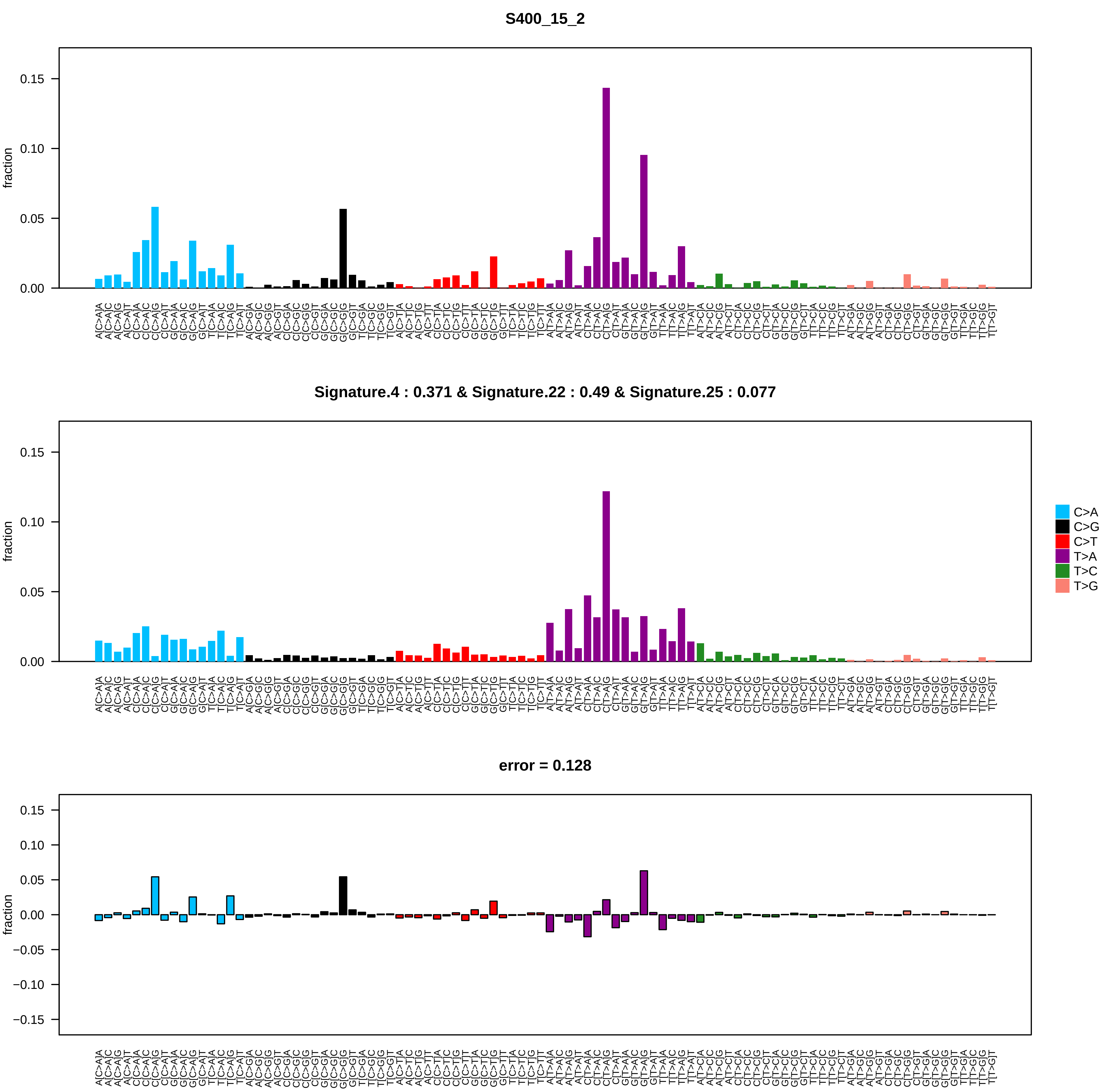

### S400_15_7.pdf

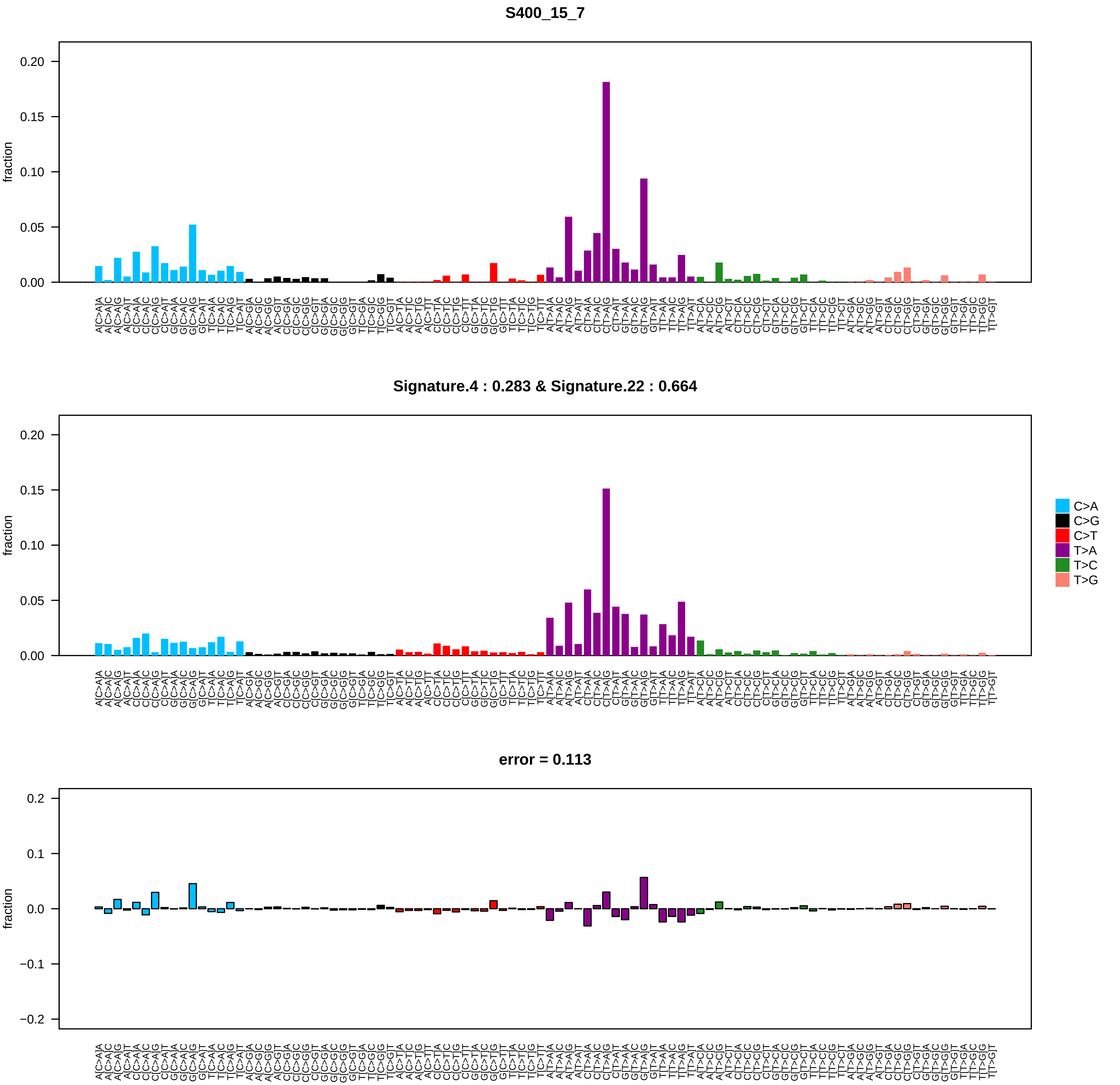

### S401_15_2.pdf

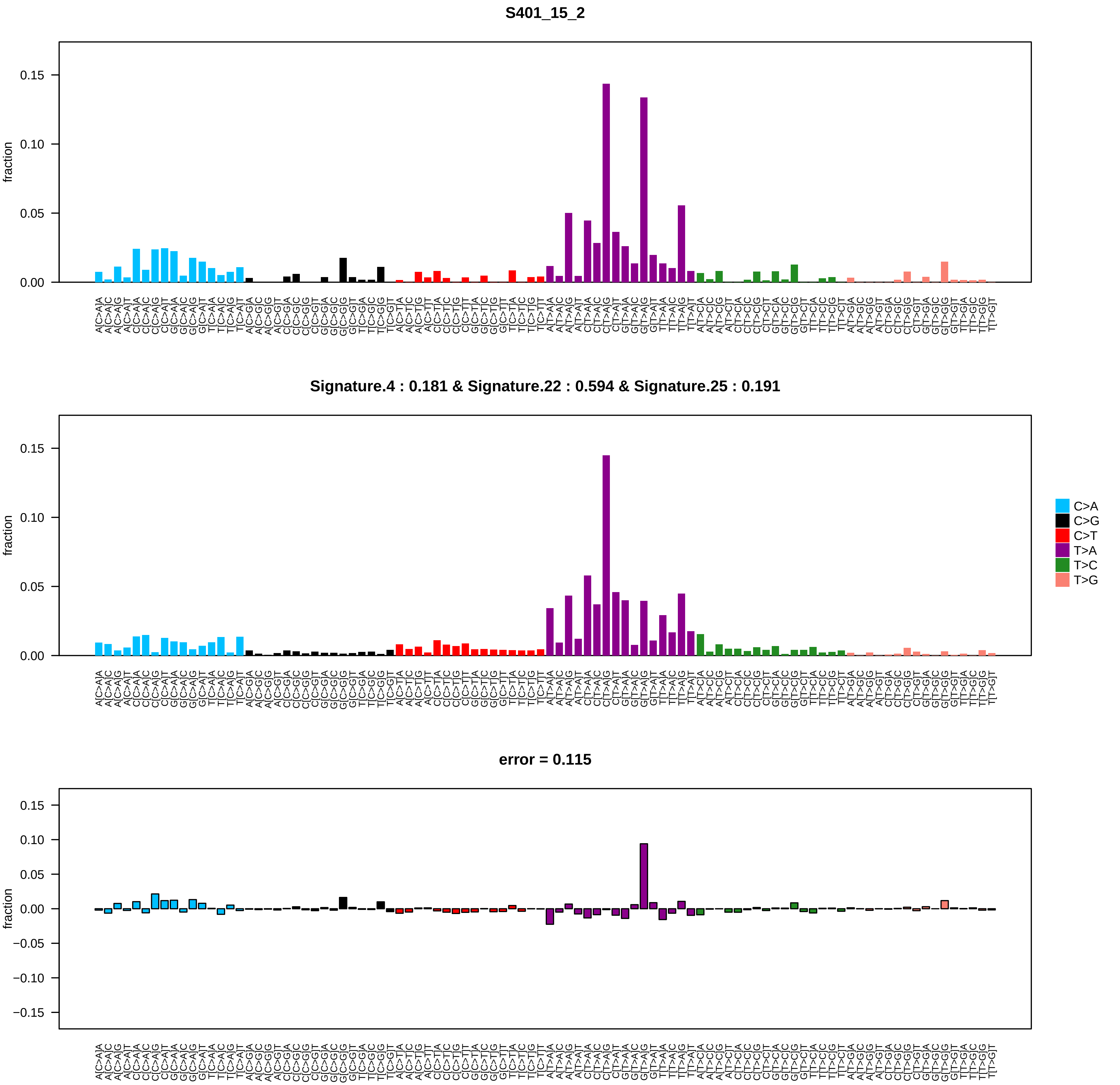

### S412_15_2.pdf

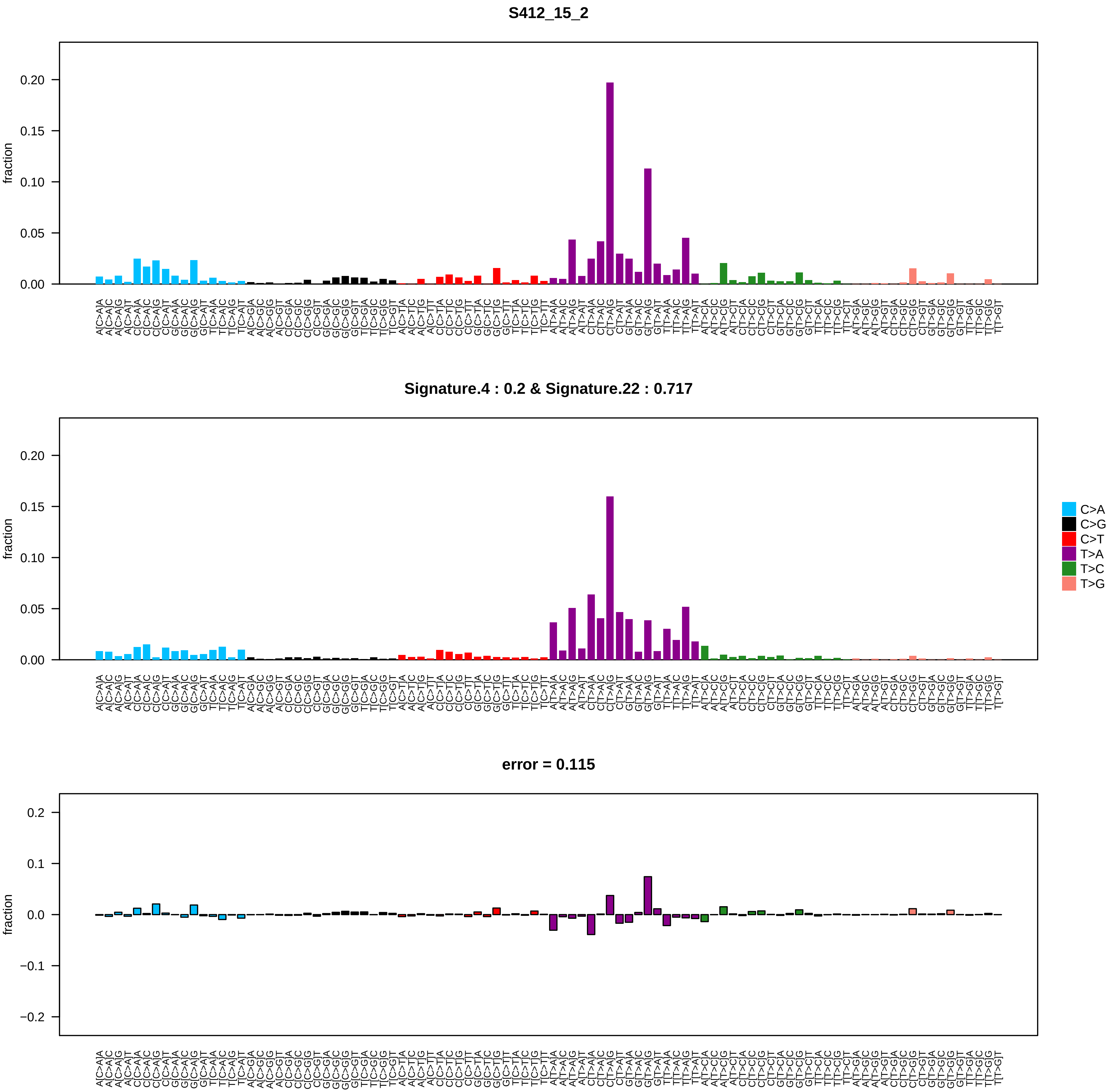

### S416_15_2.pdf

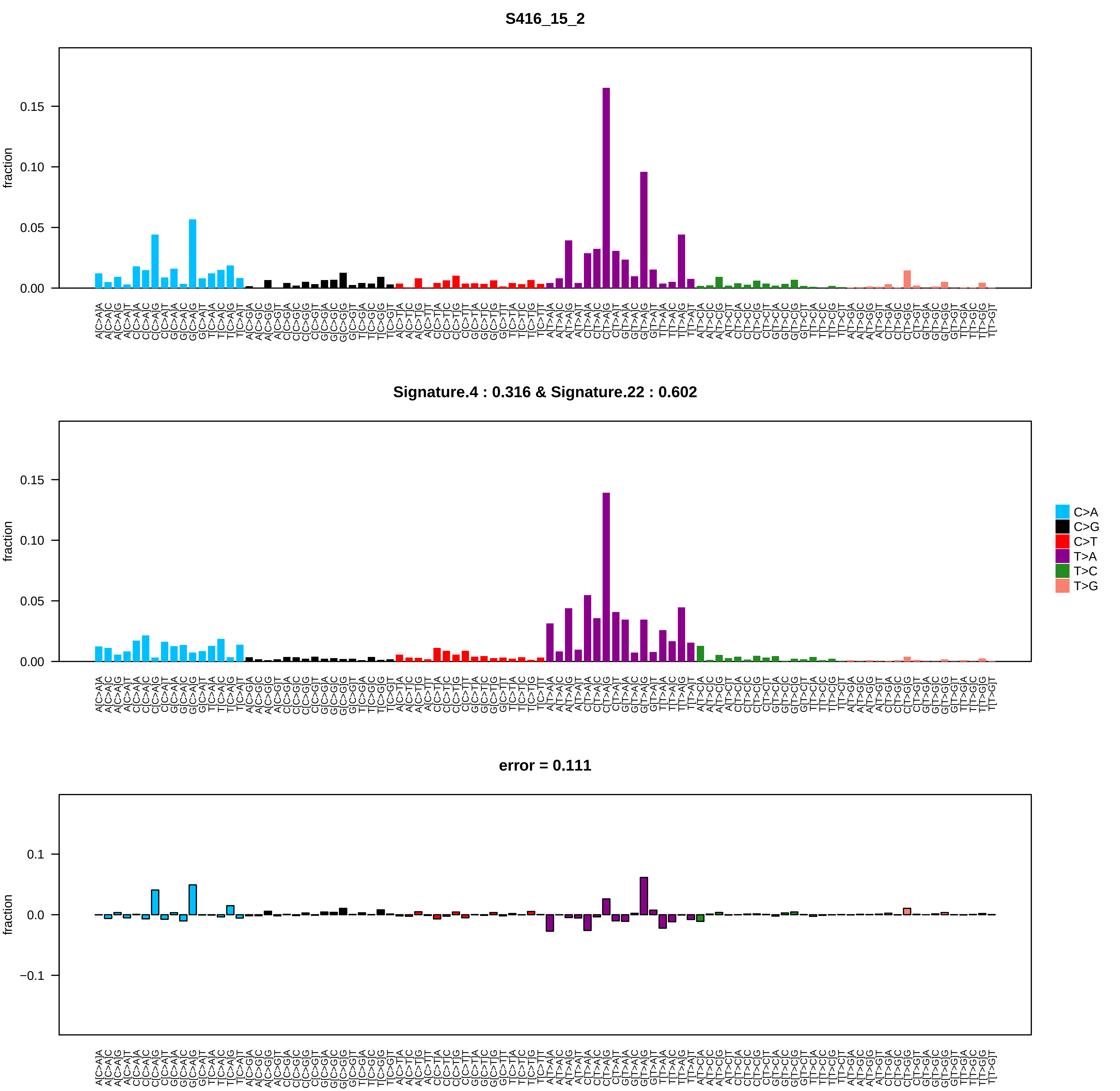

### S416_15_9.pdf

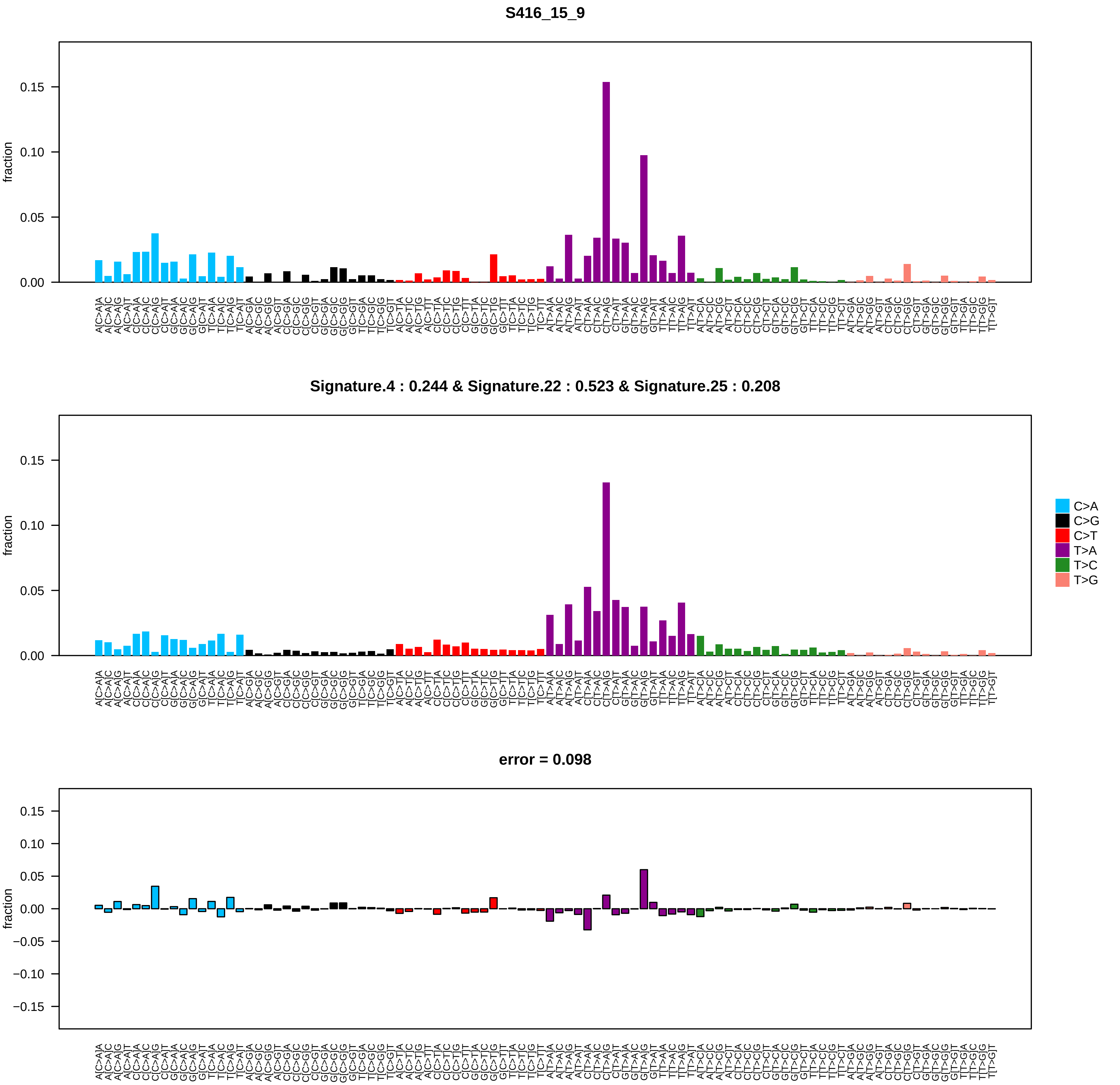

### S422_15_2.pdf

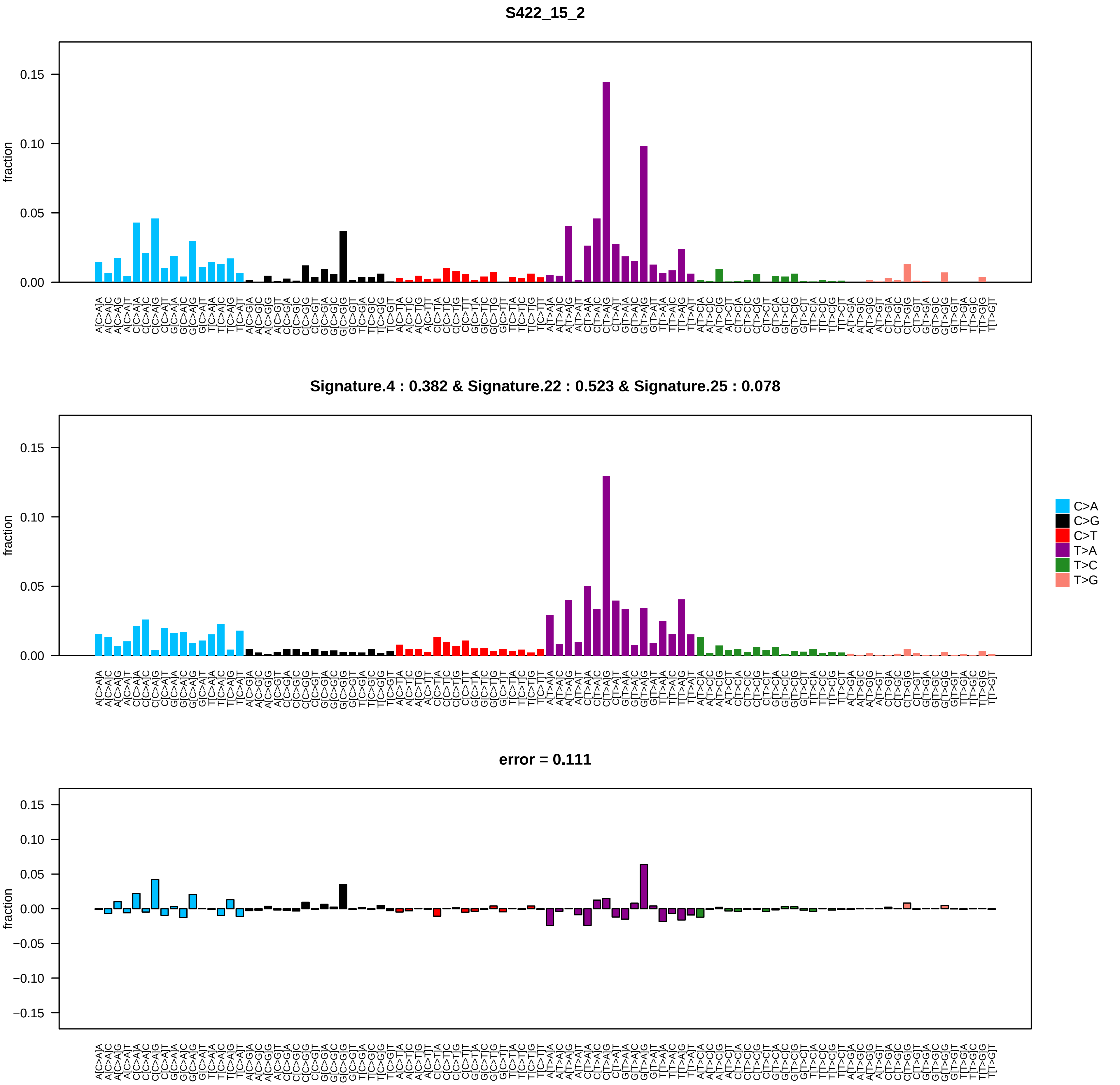
